# A molecular mechanism for membrane chaperoning by a late embryogenesis abundant protein

**DOI:** 10.1101/2022.07.29.502075

**Authors:** Xiao-Han Li, Conny W.H. Yu, Natalia Gomez-Navarro, Viktoriya Stancheva, Hongni Zhu, Cristina Guibao, Andal Murthy, Boer Xie, Michael Wozny, Benjamin Leslie, Marcin Kaminski, Ketan Malhotra, Christopher M. Johnson, Martin Blackledge, Balaji Santhanam, Douglas R. Green, Junmin Peng, Wei Liu, Jinqing Huang, Elizabeth A. Miller, Stefan M.V. Freund, M. Madan Babu

**Affiliations:** MRC Laboratory of Molecular Biology, Francis Crick Avenue, Cambridge, CB2 0QH, United Kingdom; Université Grenoble Alpes, CNRS, Commissariat à l’Energie Atomique et aux Energies Alternatives, Institut de Biologie Structurale, 38000 Grenoble, France; Department of Structural Biology and Center for Data Driven Discovery, St. Jude Children’s Research Hospital, Memphis, TN, USA; Department of Chemistry, The Hong Kong University of Science and Technology, Clear Water Bay, Hong Kong, China; State Key Laboratory of Synthetic Chemistry, Department of Chemistry, The University of Hong Kong, Pokfulam Road, Hong Kong, China; Department of Immuology, St. Jude Children’s Research Hospital, Memphis, TN, USA; Center for Proteomics and Metabolomics, St. Jude Children’s Research Hospital, Memphis, TN, USA

## Abstract

Environmental stress can result in substantial damage to proteins, membranes, and genetic material, impacting organismal survival^1-3^. Stress tolerance can be conferred by intrinsically disordered proteins (IDPs)^4^ that lack stable tertiary structure. IDPs from the large family of late embryogenesis abundant (LEA) proteins confer a fitness advantage when heterologously expressed^5,6^. Such protection suggests a general molecular function leading to stress tolerance, although the mechanisms remain unclear. Here, we report that a tardigrade LEA protein that confers stress tolerance in yeast acts as a molecular chaperone for the mitochondrial membrane. This protein, named HeLEA1, localizes to the mitochondrial matrix, and harbors conserved LEA sequence motifs that undergo dynamic disorder-to-helical transition upon binding to negatively charged membranes. Yeast expressing HeLEA1 show increased mitochondrial membrane fluidity, increased membrane potential, and enhanced tolerance to hyperosmotic stress under non-fermentative growth without significantly altering mitochondrial lipid composition or triggering a generic stress response. We demonstrate that membrane binding ameliorates excess surface tension, possibly by stabilizing lipid packing defects. Evolutionary analysis suggests that HeLEA1 homologs localize to different membrane-bound organelles and share similar sequence and biophysical features. We suggest that membrane chaperoning by LEA proteins represents a general biophysical solution that can operate across the domains of life.

## Introduction

Organisms often face environmental challenges that can affect their survival. Such challenges include extreme temperatures (heat or cold shock), water stress (desiccation or freezing) as well as altered osmotic pressure, and can result in substantial damage to proteins, membranes, and genetic material^1-3^. To survive these conditions, organisms adopt multiple strategies. Whereas large animals can physically move from harsh environments, organisms such as plants, microbes, and small animals that cannot move quickly have evolved cellular mechanisms to combat abiotic stress. One common response to diverse stresses is the expression of proteins that can increase levels of intracellular osmolytes and chemical chaperones such as trehalose, betaine, glycine^7,8^, and/or proteins that have a direct protective role on cellular components^9,10^. A subset of the latter category includes proteins with no defined tertiary structure, typically referred to as intrinsically disordered proteins (IDPs)^4^. For instance, small heat shock proteins (sHSPs) function as chaperones that protect other proteins from misfolding and aggregation^11-13^.

The late embryogenesis abundant (LEA) proteins are among the earliest IDP families discovered as combating multiple environmental stresses^5^. LEA proteins feature multiple low-complexity sequence repeats^14^, and are widely expressed in plants but also found in microorganisms and some invertebrates^5^. LEA proteins confer tolerance in the face of multiple types of stress, even when heterologously expressed^15-17^. *In vitro* experiments have suggested that LEA proteins and other IDPs adopt a vitrified state or form fibrous gels upon stress that might chaperone cellular proteins^18,19^. Additional evidence also implicates a role for IDPs in modulating vitrification of small molecules like sugars^20,21^. Moreover, LEA proteins have also been linked to broader protective functions, including enhancing function of membrane-bound organelles^22,23^, or acting as “molecular shields” to protect proteins against aggregation^24^. Consistent with a function that directly impacts membranes, *in vitro* studies have suggested that LEA proteins may protect liposomes from deformation during desiccation or freezing^25-27^. Despite extensive efforts, clear molecular functions of LEA proteins and mechanisms by which IDPs more broadly confer stress tolerance remain poorly understood. Molecular insight into such function is important in the context of the evolutionary conservation of such proteins and their roles in stress-protection.

Here we set out to define conserved mechanisms by which IDPs confer stress tolerance using a multidisciplinary approach. A combination of cellular, biophysical, structural, and evolutionary analyses suggests that a tardigrade LEA protein, named HeLEA1, chaperones mitochondrial membranes. HeLEA1 belongs to a family of LEA proteins that are predicted to localize to various membrane-bound organelles. Evolutionarily conserved LEA motifs undergo dynamic interactions with negatively charged lipids to protect the bilayer from thermal fluctuation and excess membrane tension. We propose that natural selection has preserved sequence features that drive both a disordered state and a membrane-bound structured state to confer function as a molecular chaperone for lipid bilayers in different sub-cellular organelles across diverse organisms.

### HeLEA1 is an evolutionarily conserved IDP that localizes to the mitochondria matrix

Previous work has shown that heterologous expression of tardigrade IDPs in unicellular organisms such as *Saccharomyces cerevisiae* enhances their tolerance to desiccation^18^. Inspired by this, we performed a comprehensive sequence search with HMMer^28^ to identify all remote homologs of the four tardigrade proteins demonstrated to confer desiccation tolerance when expressed in other organisms^18^. We identified 144 homologs of these four proteins, subjected them to an all-against-all sequence comparison and clustered them using the Enzyme Function Initiative similarity tool^29^. Our analysis identified two clusters: three of the four queried sequences belong to the cytosolic abundant heat soluble (CAHS) protein family, which contains only tardigrade proteins (**Extended Data Fig. 1 and Extended Data Table 1**; 48 sequences from three tardigrade species). The fourth protein belongs to a different family that is evolutionarily conserved and clusters with a group of LEA proteins (Pfam: PF02987, Group 3^30^, LEA_4^31^, or Group 6^32^ according to different classifications) (**Extended Data Fig. 1-2, Extended Data Table 2**; 96 sequences from 44 species, including two tardigrade species). We renamed this tardigrade protein, which was previously classified as a CAHS protein (UniProtID: P0CU49) as HeLEA1 (i.e., <u>LEA</u> protein from *<u>H</u>ypsibius <u>e</u>xemplaris*).

HeLEA1 homologs are found across diverse species, including bacteria, fungi, plants, and invertebrates, but not vertebrates (**Extended Data Fig.2 and Extended Data Table 2**). This distribution suggests conservation of HeLEA1 over millions of years of evolution in organisms that require molecular responses to stress. Computational analysis of these sequences suggested that different family members might localize to distinct subcellular compartments and organelles, including the nucleus, chloroplast, and mitochondria (**Fig. 1a and Extended Data Table 3**). HeLEA1 itself was predicted to target to mitochondria via a cleavable 38-residue mitochondrial-targeting sequence (MTS) at its N-terminus (**Extended Data Fig. 3**). Alignment between HeLEA1 and representative homologs revealed reasonable sequence similarity, but also long gaps (**Extended Data Fig. 4**). This is probably due to the high propensity for HeLEA1 homologs to be intrinsically disordered (83 of 96 with a mean IUPred disorder score > 0.5) (**Fig. 1b, Extended Data Fig. 5**), and the tendency for LEA proteins broadly to feature multiple copies of sequence repeats that vary in number. We performed a motif search using MEME algorithm^33^ across 96 homologs and found conserved LEA protein motifs distributed along the HeLEA1 sequence with high confidence (**Fig. 1b-Fig. 1c**). We therefore consider HeLEA1 an ideal model to investigate conserved molecular mechanisms by which LEA-related IDPs confer stress tolerance.

**Fig. 1.**
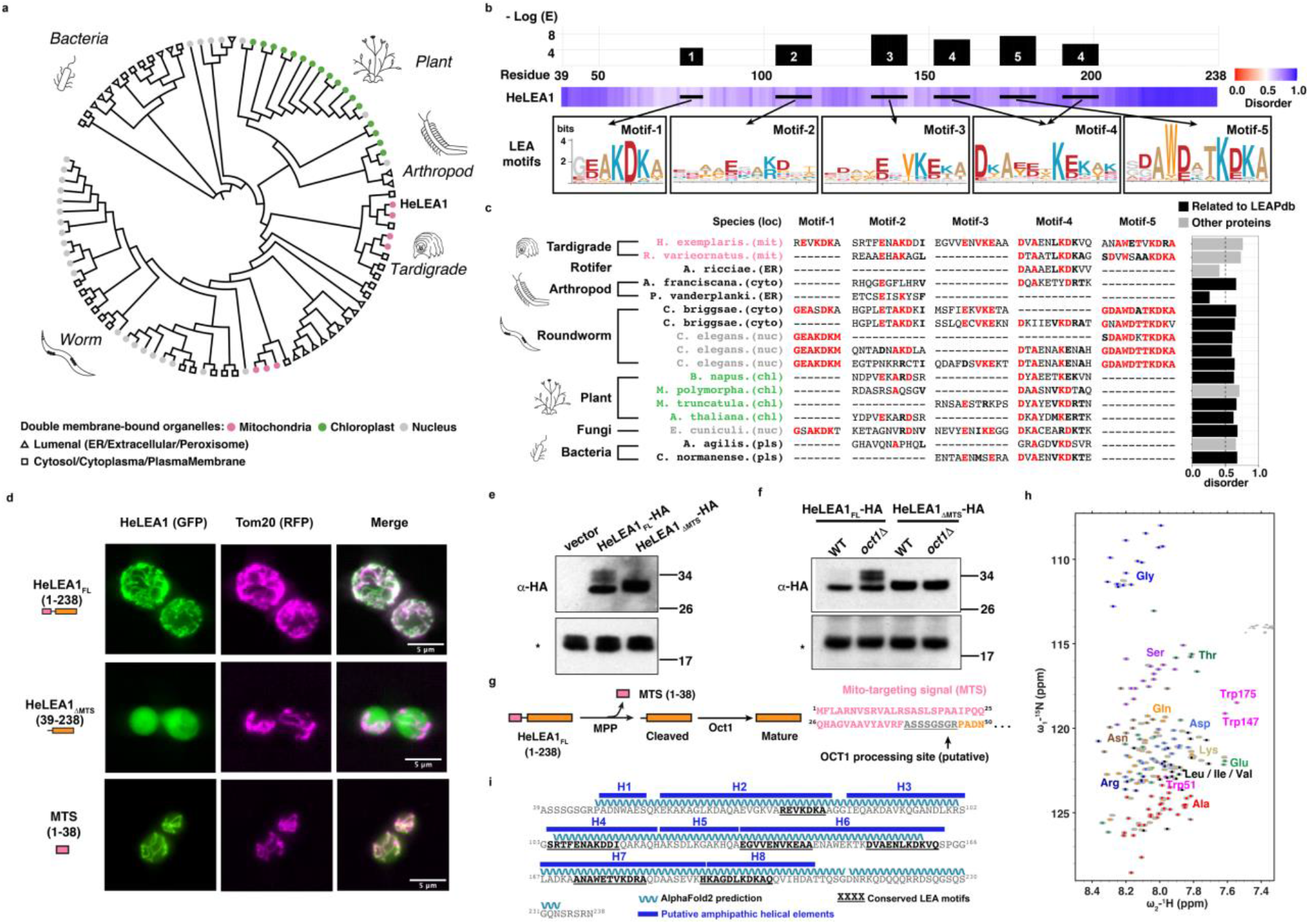
HeLEA1 is an evolutionarily conserved IDP. **a**, Evolutionary tree of HeLEA1 homologs annotated by their predicted subcellular localization. Several HeLEA1 homologs localize to various membrane-bound organelles. HeLEA1 was predicted to localize to the mitochondria. **b**, HeLEA1 possesses multiple LEA protein motifs. Top panel shows residue-specific confidence score of LEA motif in HeLEA1 (E score < 1E-4, MEME algorithm^33^). Bottom panel shows disorder prediction for HeLEA1 using IUPred2A. The locations of each high confidence motif are indicated by black lines, and the corresponding motif logos shown below. **c**, Local motif alignments between HeLEA1 and representative HeLEA1 homologs from diverse species and localizations. Mean disorder score for each homolog are stacked on the right. Homologs related to the recorded entries in the LEA protein database (LEAPdb) are highlighted in black. **d**, GFP-tagged (C-terminal) full-length HeLEA1 (HeLEA1_FL_-GFP) colocalized with mitochondria (marked by Tom20-RFP) when expressed in yeast; removal of the N-terminal 38 amino acids corresponding to the predicted MTS (HeLEA1_ΔMTS_-GFP) resulted in cytoplasmic localization; fusing the predicted MTS to GFP also resulted in mitochondrial localization (MTS-GFP). **e**, Immunoblottingimmunoblotting of log-phase cells expressing HA-tagged HeLEA1_FL_ revealed three species that likely correspond to full-length HeLEA1, MPP-cleaved HeLEA1, and mature HeLEA1 with additional processing by mitochondrial processing enzymes. HeLEA1_ΔMTS_-HA migrated as a single band of intermediate molecular weight. **f**, Immunoblotting for HeLEA1_FL_-HA and HeLEA1_ΔMTS_-HA in an *oct1*Δ background demonstrated significant accumulation of processing intermediates only for HeLEA1_FL_, supporting its mitochondrial matrix localization. **g**, Proposed protein processing events for HeLEA1 (orange): MTS (pink) is cleaved by MPP with the mitochondrial intermediate peptidase Oct1 further processing N-terminal amino acids (grey). **h**, The ^1^H_N_,^15^N 2D HSQC of HeLEA1_ΔMTS_ purified from *E. coli* shown with assignment of backbone resonances. Different types of amino acids are represented by different colours. For full assignment of the HSQC spectrum please refer to **Extended Data Fig. 8. i**, Locations of putative amphipathic helical elements predicted by biophysical properties (blue), helical regions predicted by AlphaFold2 (cyan ribbons), overlapping with conserved LEA motifs (black bold) in HeLEA1.

To test the subcellular localization of HeLEA1, we tagged full-length HeLEA1 (HeLEA1_FL_) with GFP at the C-terminus and expressed the fusion construct in *Saccharomyces cerevisiae*, which lacks HeLEA1 orthologs. HeLEA1_FL_ co-localized with a mitochondrial marker, Tom20^34^, whereas a mutant that lacked the putative MTS (HeLEA1_ΔMTS_) accumulated in the cytoplasm. Moreover, appending the predicted MTS to GFP resulted in mitochondrial localization, suggesting the predicted MTS is both necessary and sufficient for mitochondrial targeting (**Fig. 1d**). Consistent with delivery to the mitochondrial matrix, immunoblot analysis of yeast lysates from mid-log phase cells expressing HA-tagged HeLEA1_FL_ or HeLEA1_ΔMTS_ revealed multiple bands for HeLEA1_FL_, the smaller of which co-migrated with the single HeLEA1_ΔMTS_ band (**Fig. 1e**). In cells from stationary phase, HeLEA1_FL_ migrated as a single band with slightly lower apparent molecular weight than HeLEA1_ΔMTS_ (**Extended Data Fig. 6**). These observations are consistent with HeLEA1_FL_ being targeted to the mitochondrial matrix, then cleaved by the mitochondrial processing peptidase (MPP), followed by a further processing step. Indeed, when HeLEA1_FL_ was expressed in an *oct1*Δ strain that lacks a mitochondrial matrix peptidase that acts downstream of MPP^35^, more of the larger processing intermediates accumulated, suggesting matrix localized Oct1 is involved in processing of HeLEA1_FL_ (**Fig. 1f**). We conclude that HeLEA1 is a mitochondrial matrix protein and that its mature form is HeLEA1_ΔMTS_ (hereafter referred to as HeLEA1), with potential further processing at an Oct1 cleavage site (**Fig. 1g**).

To characterize the secondary structure of HeLEA1, we expressed and purified the recombinant protein from bacteria and investigated its secondary structure using circular dichroism (CD). The CD spectra of HeLEA1 showed characteristic signatures of random coil, suggesting the protein is largely disordered (**Extended Data Fig. 7**). To gather residue-specific structural information, we further characterised isotopically labelled HeLEA1 by solution-state nuclear magnetic resonance (NMR) spectroscopy. Using a combination of standard triple resonance experiments and (^1^H-start) ^13^C-detect experiments, backbone resonances for 191 out of 200 residues (90%) in HeLEA1 were assigned at 278 K (**Fig. 1h, Extended Data Fig. 8**). Two features stood out in the ^1^H_N_-^15^N 2D HSQC spectra: (a) very narrow dispersion of amide proton chemical shifts, and (b) clustering of the backbone ^1^H_N_,^15^N resonances in the 2D spectrum according to amino acid type. These features define HeLEA1 as an intrinsically disordered protein in solution.^36^

In addition to the general feature of predicted disorder in solution (**Extended Data Fig. 5**) and conserved LEA motifs, our sequence search suggested that some HeLEA1 homologs harbour sequence motifs with similarity to the apolipoprotein superfamily (Pfam) (**Extended Data Fig. 9**). Additionally, a PhyRE2^37^ structural similarity search probing the PDB database with the HeLEA1 sequence yielded hits with apolipophorin-III, α-synuclein and HSP12 (**Extended Data Fig. 10**), all of which associate with membranes via amphipathic helices^38^. We therefore devised a computational pipeline to identify potential 3-11 helical elements (minimum length = 7 residues, with aligned amphipathic interface) that may interact with lipids according to previously reported criteria for predicting continuous amphipathic helical elements^39^ and apolipoprotein motifs^40,41^ (**Extended Data Fig. 11**). We identified eight putative amphipathic 3-11 helical elements of various lengths in HeLEA1 (H1–H8, **Fig. 1i, Extended Data Fig. 12**). These elements are enriched in weakly hydrophobic residues (alanines and threonines) and exhibit a small hydrophobic surface (3–5 of the 11 projected positions is hydrophobic). Moreover, four of the eight helices (H2, H6–H8) displayed strong enrichment of positively charged residues (lysines and arginines) at the boundary of the hydrophobic and hydrophilic surfaces, and enrichment of negatively charged residues (glutamates and aspartates) in the hydrophilic surface. The other helices (H1, H3–H5) also exhibit these features, albeit weakly. The biophysical prediction corresponds well with PSIPRED (**Extended Data Fig. 13**) and AlphaFold2 prediction^42^ (**Extended Data Fig. 12, Fig. 1i**). The low pLDDT score of the AlphaFold2 prediction suggests a low confidence global structure (**Extended Data Fig. 12**), probably due to the disordered nature of HeLEA1 causing difficulty with sequence alignment (**Extended Data Fig. 4, Extended Data Fig. 5**). Nonetheless, local secondary structural elements predicted by AlphaFold match our biophysical prediction (**Extended Data Fig. 12**). The low confidence in stably-folded long amphipathic helices but agreement on local secondary structure predictions would be consistent with a dynamic disorder-to-helical transition in HeLEA1, a common feature of IDPs binding to membranes^43-46^ that is also shared by some LEA proteins^6^.

### Conserved LEA motifs remain conformationally dynamic during the early stages of disorder-to-helical transition of HeLEA

IDPs are frequently proposed to function as an entropic buffer, maintaining a degree of disorder even when adopting a functionally relevant conformational state^4^. Our computational analysis of HeLEA1 suggests specific sequence elements that can adopt both disordered and structurally defined states, largely consisting of amphipathic elements (**Fig. 1i**). To gain structural insight into these elements, we sought to probe HeLEA1’s conformational states and backbone dynamics in solution, using trifluoroethanol (TFE) perturbation. TFE is frequently used in protein folding studies to mimic conditions under which transient helical components of the conformational ensemble are stabilized^47^. Circular dichroism (CD) revealed that TFE indeed induced helicity of HeLEA1 in a concentration-dependent manner (**Fig. 2a**). We subsequently performed integrative structural analysis of HeLEA1 in either 0% or 10% TFE, conditions under which HeLEA1 remained predominantly disordered (**Fig. 2a inset**). Any observed residue-specific differences for these two conformational states should provide insight into disorder-to-helical transitions for each amphipathic element. In general, the biophysical properties of HeLEA1 were experimentally investigated by SAXS and NMR, and the *ASTEROIDS* algorithm^48,49^ was used to determine the conformational ensemble that best described all experimental data (**Extended Data Fig. 14**). We found that 10% TFE mildly increased the global helical propensity, with certain regions of H1 (51-54), H2 (57-62), H4 (114-120), H7 (183-187) exhibiting a significant increase in helical conformation (**Fig. 2b, Extended Data Fig. 15**). The radii of gyration (R_g_) of HeLEA1 in 0% TFE and 10% TFE were larger than the R_g_ predicted for a statistical random coil (**Fig. 2c**) and adding 10% TFE resulted in a small decrease of R_g_ compared to the 0% TFE condition (**Fig. 2c, Extended Data Table 4, Extended Data Fig. 16**). These results correspond well with HeLEA1 occupying a largely expanded disordered state, but with significant increase in helical propensity within short local segments at the early stage of disorder-to-helical transition (**Fig. 2d**).

**Fig. 2.**
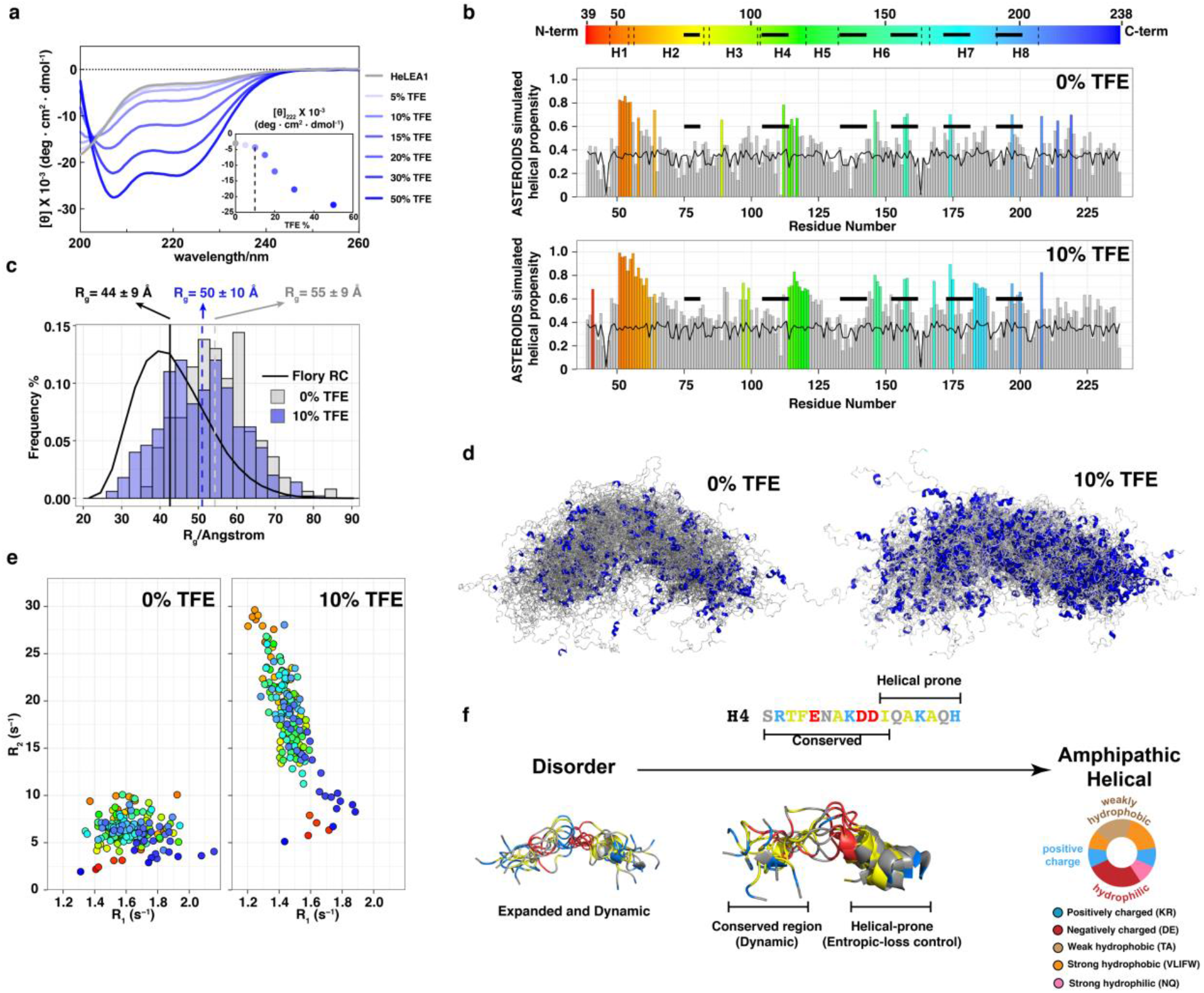
Conserved elements of HeLEA1 remain dynamic during early stages of disorder-to-helical transition. **a**, Circular Dichroism (CD) Titration of 10*µ*M HeLEA1 with trifluoroethanol (TFE). Inset shows the increase in helicity (monitored as molecular ellipticity at 222nm) as a result of increased TFE concentrations. Note that at 10% TFE (dashed line), HeLEA1 appears predominantly disordered. **b**, Residue-specific helical propensity based on *ASTEROIDS* fits of NMR data of HeLEA1 in 0% TFE (top) or 10% TFE (bottom). Residues with significant helical propensity (deviating > 0.3 from random coil (black line)) are highlighted according to the colour scheme on top. Black bars represent conserved LEA motifs mapped in **Fig. 1b. c**, Distribution of the radius of gyration (R_g_) of the HeLEA1 ensemble generated assuming random coil (RC) behaviour (black line) or fitted with SAXS data either from 0% TFE (gray bars) or 10% TFE (blue bars). **d**, Full description of the conformational ensemble (100 representative conformers) of HeLEA1 in 0% TFE (left) and 10% TFE (right) by integrating NMR and SAXS data within the *ASTEROIDS* simulation pipeline. Helical regions in each conformer are highlighted in blue. **e**, Correlation maps of transverse (R_2_) and longitudinal (R_1_) relaxation rates of HeLEA1 in 0% (left) or 10% TFE. 10% TFE drastically increased the R_2_ relaxation rates in regions with increased helical propensity, with same colour scheme as shown in **Fig. 2b. f**. Schematic depicting the disorder-to-helical transition for HeLEA1 H4 as an exemplar: weak amphipathic elements comprise a region of higher disorder propensity adjacent to a region with an increase in intrinsic helical propensity. This arrangement may reduce the cost of conformational entropy during the disorder-to-helical transition. 10 conformers for the H4 region as predicted from *ASTEROIDS* ensembles are shown.

We next probed protein backbone dynamics by ^15^N NMR relaxation measurements as HeLEA1 transits from disorder-to-helical conformation. ^15^N{^1^H} heteronuclear nuclear Overhauser effect (hetNOE) experiments probe fast (picoseconds) backbone dynamics but revealed only a small global increase in 10% TFE, potentially indicating a modest increase in overall rigidity (**Extended Data Fig. 17**). Accordingly, a modest decrease was observed for ^15^N R_1_ (1/T_1_) longitudinal relaxation rates (**Fig. 2e**). In contrast, ^15^N R_2_ (1/T_2_) transverse relaxation rates displayed a dramatic increase under 10% TFE conditions (**Fig. 2e**). ^15^N R_2_ rates sample backbone dynamics at multiple time scales and are exquisitely sensitive to ‘segmental motions’ associated with the formation of transiently populated secondary structural elements. Sequence elements with significantly increased ^15^N R_2_ rates correlate with regions exhibiting increased helicity (**Fig. 2b**). These results support the model that a disorder-to-helical transition is initiated via the population of local secondary structure elements. This results in a global decrease in backbone flexibility and shifts the overall conformational ensemble towards higher helical propensity. Importantly, the regions that mapped to evolutionarily conserved LEA motifs did not completely overlap with regions of higher helical propensity (**Fig. 2b**). The localizations of these conformationally dynamic motifs imply that maintaining a high conformational-entropy state in the conserved regions may be important in the function of LEA proteins, while the neighbouring helical-prone regions may buffer the cost of conformational entropy during the disorder-to-helical transition. (**Fig. 2f**).

### HeLEA1 stabilizes negatively charged membranes through dynamic disorder-helical transition

Our cellular localization and conformational dynamics data suggest that in its physiological setting, HeLEA1 undergoes a disorder-to-helical conformational change upon binding to the inner mitochondrial membrane (IMM). We therefore directly tested whether HeLEA1 can be recruited to membranes using a liposome flotation assay. Since the IMM is enriched in negatively charged lipids, and the amphipathic elements in HeLEA1 are enriched in positively charged residues (**Extended Data Fig. 12, Fig. 2f**), we tested HeLEA1 recruitment to small unilamellar vesicles (SUVs) made with either negatively charged POPS or neutral POPC. At physiological salt concentrations, HeLEA1 stably associated with POPS, but not POPC SUVs (**Fig. 3a**). Increasing the salt concentration, which suppresses electrostatic interactions but encourages hydrophobic interactions, abolished binding of HeLEA1 to negatively charged liposomes and promoted weak association with neutral liposomes (**Fig. 3a**). Thus, HeLEA1 can bind negatively charged membranes primarily through electrostatic interactions but may interact weakly with neutral lipids via hydrophobic interactions.

**Fig. 3.**
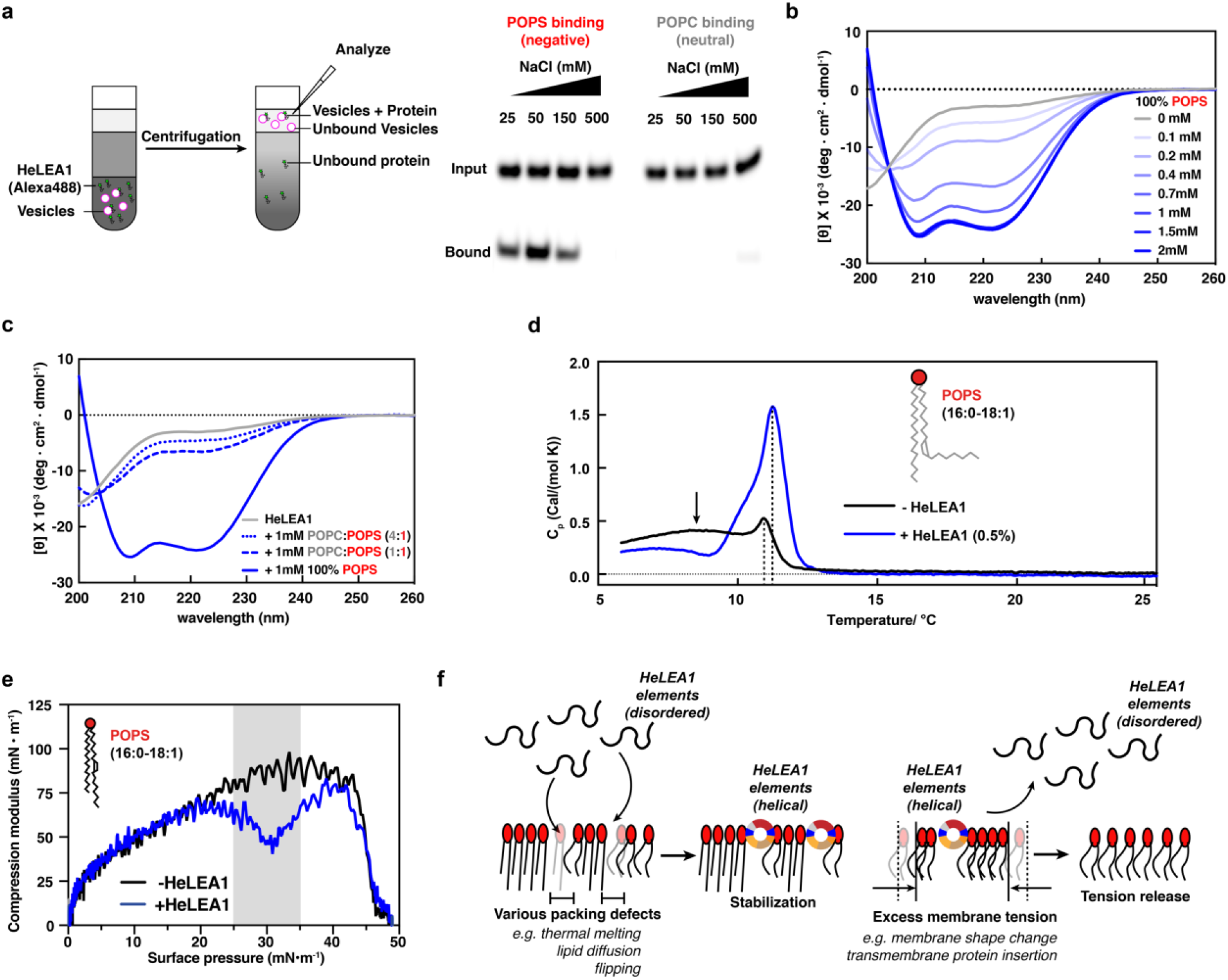
HeLEA1 stabilizes negatively charged membranes through dynamic disorder-to-helical transition. **a**, Lipid flotation assay with Alexa Fluor 488–labelled HeLEA1 and liposomes made of various lipids (POPS, red; POPC, gray) at different salt concentrations. **b**, Titration of 10µM HeLEA1 with increasing amounts of 100% POPS SUVs monitored by CD. **c**, Titration of 10µM HeLEA1 with 1mM SUVs of different ratios of negatively charged POPS versus neutral POPC, monitored by CD. Decreasing the amount of POPS fraction significantly reduced the induced helicity of HeLEA1. **d**, Differential scanning calorimetry for POPS SUVs without (black line) and with HeLEA1 (0.5% molar ratio, blue line). HeLEA1 suppressed thermally induced phase transition at low temperature (broad peaks indicated by arrow) and increased phase transition temperature (peak position; dashed lines). **e**, Change of *C*_*S*_^−1^ values in POPS monolayers with respect to surface pressure, as measured by Langmuir-monolayer methodology, with or without 3nM HeLEA1 (1:50 protein:lipid ratio). Gray areas depict the range of physiological surface pressure. **f**, Proposed mechanism for the molecular function of HeLEA1. A disorder-to-helical transition occurs within putative amphipathic elements in HeLEA1 when HeLEA1 interacts with negatively charged lipids at areas of the membrane with lipid packing defects, resulting in stabilization (left). When excess surface tension occurs at a HeLEA1-bound membrane, HeLEA1 elements would unfold and detach from the membrane to buffer such tension, stabilizing the membrane and increasing membrane fluidity (right).

We next determined whether membrane binding induces structural changes similar to those observed with TFE. We titrated HeLEA1 with SUVs containing various ratios of neutral lipids to negatively charged lipids and monitored the changes in protein secondary structure by CD. We observed increased helicity of HeLEA1 with increasing concentrations of negatively charged SUVs (**Fig. 3b**) and proportional to the fraction of negatively charged lipids (**Fig. 3c**). The CD spectra of the POPS titration revealed an isodichroic point at 204 nm, which is characteristic of a classic disorder-to-helical transition^50^. Deconvolution of the CD titration curves with Bestsel^51^ suggested that at least 60% of the residues in lipid-bound HeLEA1 are in a helical conformation, corresponding well with predictions of possible secondary structures and TFE-induced disorder-to-helical transition (**Fig. 1i, Fig. 2a**). Moreover, we observed complete reversibility of such membrane-induced disorder-to-helical transition without cooperativity in thermal melting and refolding experiments (**Extended Data Fig. 18**). Such reversibility corresponds well with the highly dynamic behaviour of HeLEA1 during its structural transition as mapped by TFE titration. Importantly, the positional distribution of the conserved dynamic LEA motifs correlates with a weak local hydrophobic moment and the presence of negative charge clusters, featuring a low lipid discrimination factor and indicating weak protein-lipid interaction (**Extended Data Fig. 19**). These features are conserved in amphipathic elements in HeLEA1 homologs, compared to similar amphipathic elements observed in apolipoproteins that is known to stably bind to lipids for structural scaffolding (**Extended Data Fig. 20**). These observations highlight the importance of weak and dynamic lipid binding in the functioning of HeLEA1.

We therefore postulated that weak binding of HeLEA1 to negatively charged membranes confers chaperone-like activity to modulate the biophysical properties of the bilayer. We first tested this model by investigating the temperature-dependent lipid phase transition of negatively charged synthetic membranes using differential scanning calorimetry (DSC) in the presence or absence of HeLEA1. DSC measures heat flux towards the sample as a result of changing temperature. A lipid bilayer will have a characteristic DSC peak triggered by phase transition from an ordered phase to a liquid disordered phase as packing of fatty acids becomes disordered with increasing temperature. In the absence of HeLEA1, POPS SUVs displayed a somewhat complex phase transition profile over a broad temperature range (**Fig. 3d**), as previously reported^11^. Addition of sub-stoichiometric amounts of HeLEA1 (1:200 molar ratio of protein:POPS) suppressed phase transition at low temperatures (arrow in **Fig. 3d**) and increased the dominant phase transition temperature (DSC peak position) (dashed lines in **Fig. 3d**). Experiments using SUVs composed of 100% DMPS, a lipid with saturated fatty acid chains, yielded similar results, whereas the behaviour of SUVs made of neutral lipids remained mostly unchanged in the presence of HeLEA1 (**Extended Data Fig. 21**). These results suggest that HeLEA1 can stabilize membranes from thermally induced fluctuations of bilayers independent of the saturation state of lipid side chains.

We next tested if HeLEA1 can also modulate mechanical properties of membranes using Langmuir-monolayer methodology^52^, in which the relationship between the surface pressure and area of a lipid monolayer is determined as compression isotherms (**Extended Data Fig. 22**). The resulting curves are used to compute the compression modulus (*C*_*S*_^−1^), which reports on the stiffness of the lipid monolayer: lower *C*_*S*_^−1^ values correspond to softer and more compressible membranes with increased fluidity^53^. The presence of 3nM HeLEA1, which translates to a protein-to-lipid ratio of 1:50, substantially decreased *C*_*S*_^−1^ (i.e., increased membrane fluidity) for POPS monolayers (**Fig. 3e**) but marginally changed the *C*_*S*_^−1^ for POPC monolayers (**Extended Data Fig. 23**), in good agreement with its preferred recruitment to negatively charged phospholipids. Notably, this effect was most prominent in the physiological range of surface pressures, i.e., 25–35 mN/m (**Fig. 3e**, gray box)^53^. The mean *C*_*S*_^−1^ of the POPS monolayer changed from 85.4±5.8 mN/m to 54.8±6.6 mN/m at physiological membrane surface pressure, suggesting a significant increase in membrane fluidity^53^. At high surface pressure, the *C*_*S*_^−1^ of the POPS monolayer with HeLEA1 was restored to that without HeLEA1, suggesting that HeLEA1 fall out of the lipid monolayer, suggesting a reversible and dynamic protein-lipid interaction.

Together, our biochemical data with synthetic bilayers are consistent with a role for HeLEA1 as a conserved molecular chaperone for lipid bilayers that promotes stabilization and modulation of membrane fluidity. The evolutionarily conserved dynamic disorder-helical transition features in HeLEA1 contribute to its chaperone function (**Fig. 3f**).

### HeLEA1 expression enhances tolerance to hyperosmotic stress on non-fermentable carbon sources

To further explore HeLEA1 activity under more physiological conditions, we first tested recruitment to giant unilamellar vesicles (GUVs) that mimic the IMM composition^54^. We encapsulated N-terminally fluorescently labelled HeLEA1 inside the GUVs and imaged both lipid and protein by confocal imaging. HeLEA1 colocalized with the GUV membrane (**Fig. 4a**), suggesting that HeLEA1 binds membranes of a physiologically relevant composition. We next aimed to observe HeLEA1’s function on mitochondria more directly. We generated a yeast strain that expresses HeLEA1_FL_ under a strong constitutive promoter, purified mitochondria and stained them with diphenylhexatriene (DPH). DPH fluoresces in a hydrophobic environment, and its anisotropy reports on membrane fluidity, where low anisotropy (faster tumbling) corresponds to a more fluid membrane (less ordered lipids). Mitochondria from HeLEA1-expressing cells showed decreased DPH anisotropy, i.e., increased membrane fluidity, compared to wild type cells (**Fig. 4b**). This change in fluidity is unlikely to be driven by changes to the lipid composition because principal component analysis of lipidomics experiments from isolated mitochondria revealed no significant difference in composition between these two conditions (**Extended Data Fig. 24**).

**Fig. 4.**
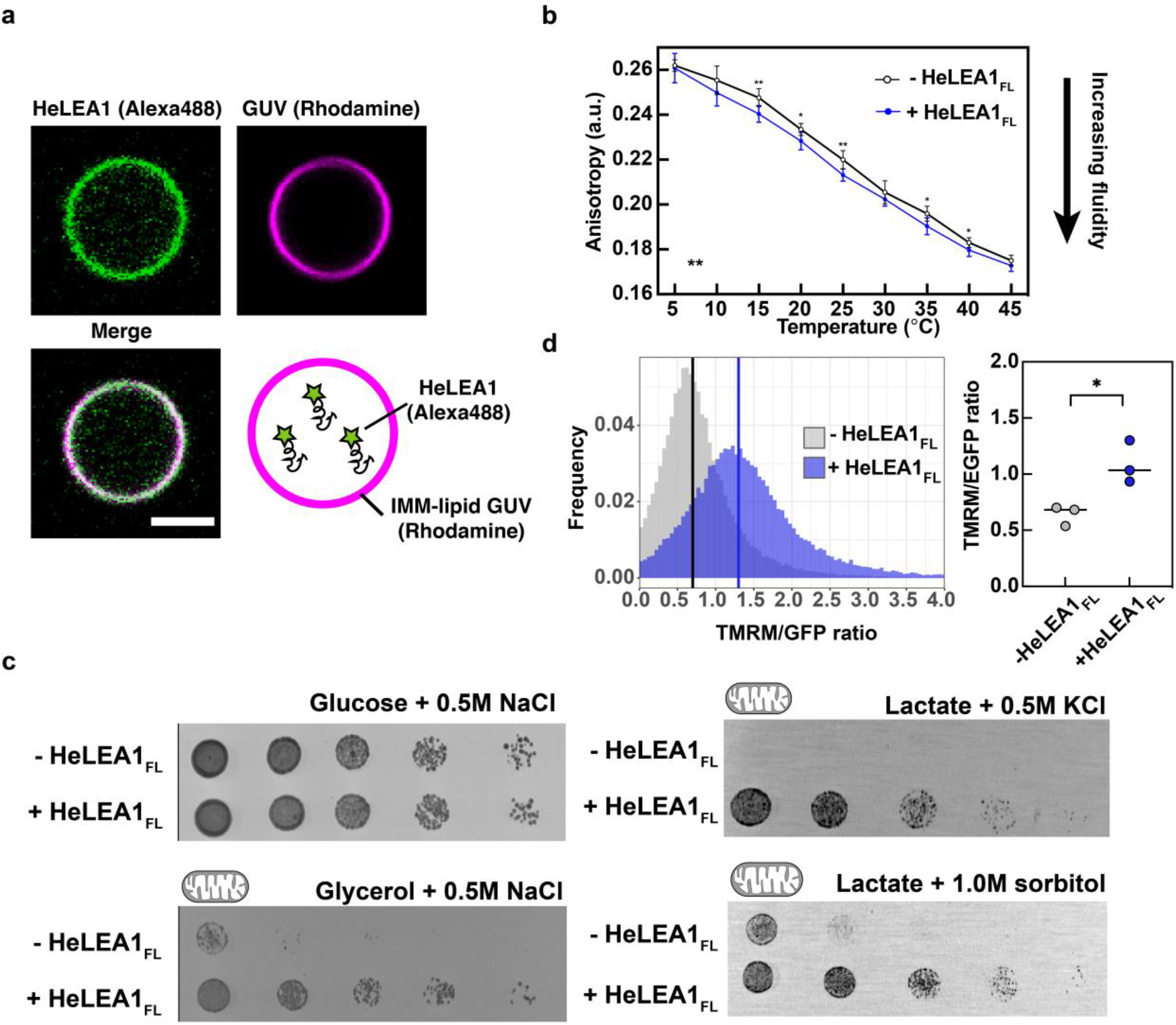
Physiological impact of HeLEA1 expression. **a**, Confocal imaging showing colocalization of HeLEA1 and GUVs with a composition mimicking that of the inner mitochondrial membrane (IMM). GUVs were labelled with DOPE-Lissamine rhodamine (magenta). HeLEA1 was labelled with Alexa Fluor 488 (green) and encapsulated inside GUVs. Scale bar: 3 µm. **b**, Temperature-dependent fluorescence anisotropy of DPH-stained mitochondria from yeast cells grown in normal YPD media. Mitochondria with HeLEA1_FL_ expression had lower anisotropy than without HeLEA1_FL_ expression (**, *P* < 0.01, two-way ANOVA, *n* = 7, error bars represent the standard deviation), suggesting that HeLEA1_FL_ expression increases the membrane fluidity of mitochondria. For each temperature pairs, an independent unpaired t-test were also performed, and the statistical significance was indicated (*, *P* < 0.05, **, P < 0.01, two-tailed, unpaired t-test, *n* = 7). **c**, HeLEA1_FL_ expression had a positive fitness effect on various hyperosmotic stress when cells were grown on non-fermentable carbon sources, in which mitochondrial activity is crucial for cell survival, but not with fermentable carbon source glucose where mitochondrial activity is less critical for survival. **d**, Normalized TMRM fluorescence for yeast cells with or without HeLEA1_FL_ expression grown on glycerol. HeLEA1_FL_ expression significantly increase mitochondrial membrane potential, suggesting enhanced mitochondrial activity in live cells. Representative FACS data for one replica (left, lines indicate median of the distribution) and statistical test for three independent replica (*, *P* < 0.05, Welch t-test, *n* = 3, right) are shown.

Hyperosmotic stress is commonly associated with upregulation of stress-related IDPs including LEA proteins^2,55,56^, and altered mitochondrial function^57,58^. We therefore tested if HeLEA1 could confer enhanced tolerance to salt stress (0.5M NaCl). Strikingly, HeLEA1_FL_ expression conferred a substantial growth advantage only when grown on non-fermentable glycerol media at 37 °C, where cell growth is dependent on mitochondrial function^59^. In contrast, on glucose media, where mitochondrial biogenesis is repressed and mitochondrial function is not essential, there was no growth advantage (**Fig. 4c**). We observed similar effects in cells grown on lactate, another non-fermentable carbon source, and using other hyperosmotic stress conditions including 0.5 M KCl and 1.0 M sorbitol (**Fig. 4c, Extended Data Fig. 25**). Similar phenotypes were also observed when HeLEA1_FL_ was expressed under a weaker promoter (**Extended Data Fig. 26**). Overexpression of GFP with the HeLEA1 MTS did not confer stress protection to the same extent (**Extended Data Fig. 27**), suggesting the phenotype is not caused by mitochondrial import stress. Similarly, altered lipid composition of the mitochondria is unlikely to explain the growth phenotypes since we did not observe significant changes in mitochondrial lipid composition in the presence or absence of HeLEA1_FL_ under glycerol growth (**Extended Data Fig. 28**). Instead, we observed that HeLEA1_FL_ expression conferred a higher mitochondrial membrane potential. For this, we stained mid-log phase cells growing in glycerol with TMRM, a fluorescent dye whose accumulation in mitochondria is dependent on membrane potential. We normalized TMRM fluorescence to mitochondrial abundance using TOM20-eGFP. HeLEA1_FL_ expression resulted in higher TMRM fluorescence consistent with increased mitochondrial membrane potential (**Fig. 4d**). As HeLEA1 does not have any homologs identified in yeast (**Extended Data Table 2**), these phenotypes because of heterologous expression of HeLEA1 strongly support that membrane chaperoning by HeLEA1 promotes mitochondrial function may serve as a general biophysical mechanism for stress tolerance.

## Discussion

Since the discovery of LEA proteins from cotton seeds, the relevance of these IDPs for stress tolerance has been demonstrated in multiple organisms and model systems, including tardigrades^23,60^ and arthropods^12,15,61^. However, the molecular mechanism by which these stress-tolerance proteins exert their protective function has remained less clear. Using the tardigrade HeLEA1 protein as a model, we performed a detailed characterization at multiple scales (atomic, molecular, subcellular, cellular, and evolutionary) to uncover a possible general molecular mechanism by which the IDPs of the LEA family confer stress tolerance.

We showed that HeLEA1 is an IDP that localizes to mitochondria and undergoes a disorder-to-helical transition in the presence of negatively charged lipids. The high conformational entropy of the intrinsically disordered state stabilizes HeLEA1 in solution, whereas HeLEA1 folding upon binding to negatively charged lipids trades a loss of conformational entropy for enthalpic gain from electrostatic interactions, exposing the hydrophobic surfaces of the weakly amphipathic helical elements. This mechanism explains the importance of the conserved LEA motifs (**Fig. 2f**), which confer reversible association with lipids. Such dynamic interactions may buffer local changes in the surface tension and packing defects of membranes (**Fig. 3f**). Because the IMM is enriched in negatively charged lipids, including cardiolipin^62,63^, the dynamic membrane-binding process of HeLEA1 may thereby encourage the diffusion of transmembrane proteins^64^ in the IMM as well as buffering changes in membrane tension during stress and membrane protein biogenesis. The notion is further supported by evidence where increased membrane fluidity of mitochondria by altering unsaturation side chains enhance their function^65^.

Our observations that HeLEA1 only confers significant hyperosmotic stress tolerance on non-fermentable carbon source, when mitochondrial function is essential, suggest that the molecular function of an IDP (i.e., its biophysical properties and behaviour) can be robust, while its cellular function (i.e., effect on mitochondrial activity) is context dependent. Our evolutionary analysis suggests that amphipathic elements in HeLEA1 homologs with different predicted targeting signals display trade-offs between fraction of basic residues and hydrophobicity, while keeping the hydrophobic moments unchanged (**Extended Data Fig. 30**). We propose that the molecular function of weak lipid binding (i.e. low hydrophobic moments) is conserved, but the mechanism for lipid recognition (electrostatic-driven versus hydrophobic-driven) is fine-tuned according to diverse subcellular localizations, where different membrane-bound organelles differ in their lipid compositions^66,67^. Proteins with distant structural similarity to HeLEA1, such as apolipoproteins, may have evolved away from the need to protect organelles from environmental stresses and acquired new functions, such as stabilization of lipoprotein particles in mammals. Accordingly, their sequences evolved to change their biophysical properties from dynamic and disordered to stable and folded (**Extended Data Fig. 19**).

The expression of a tardigrade protein that confers stress tolerance in yeast highlights the generic biophysical basis for stress tolerance through functional enhancement of membrane-bound organelles. Therefore, HeLEA1 homologs may confer abiotic stress tolerance by acting as membrane chaperones, i.e., maintaining the membrane fluidity of membrane-bound organelles and facilitating proper biological functions of their associated membrane proteins (**Fig. 5**). We suggest that such a biophysical mechanism may represent a simple and evolvable solution that can confer stress tolerance across the different domains of life.

**Fig. 5.**
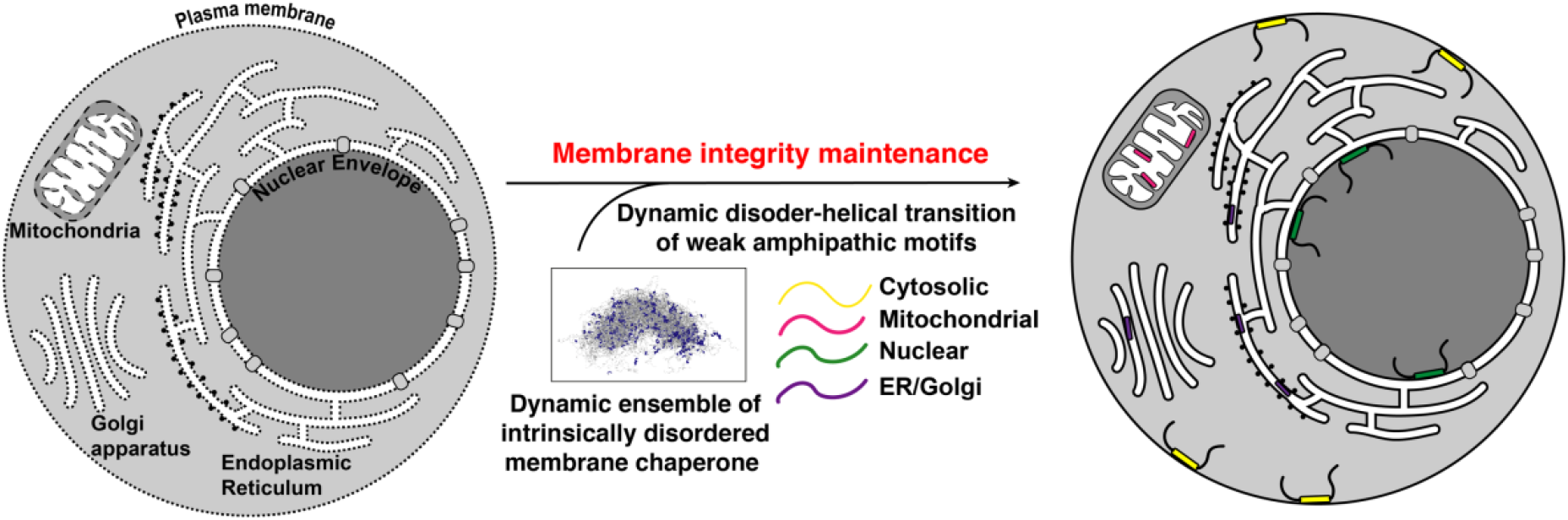
Proposed function for HeLEA1 homologs. HeLEA1 homologs may confer various stress tolerance phenotype by acting as membrane chaperones targeted to different subcellular localizations.

## Acknowledgments

We thank A. Gunnarson, E. Rhoades, R. Kriwacki, I. Chen, A. Elazar, and B. Lang for reading and critical comments on the manuscript; A. Carter and C. Lau for helping with protein purification; E. Derivery, J. Watson and H. McMahon for helping with GUV construction; and Y. Ohashi for kindly providing the yeast strain for imaging. We thank T. Boothby for providing the original plasmid hosting tardigrade proteins. We thank Z. Chen, J. Lu, G. Slodkowicz, Z. Shi, Y. Yagita for helpful discussions on GUV construction, statistical analysis and interpretation of the data. We also thank Diamond Light Source for Beamtime (proposal SM24985) and the staff of Beamlines B21 for assistance with SAXS data collection; H. Tan, J.-H. Cho, and St. Jude Center for Proteomics and Metabolomics for assistance in lipidome profiling; S. Mclaughlin and LMB Biophysics Instrument Centre for assistance in CD and fluorescence data collection; J. Howe and the LMB light microscopy team for assistance in microscopy; M. Daly and the LMB cell sorting facility for assistance in flow cytometry; LMB Scientific Computing for providing computational resources for simulation; and LMB media and glass wash for helping to prepare media and plates. This work was supported by the Medical Research Council, as part of United Kingdom Research and Innovation (also known as UK Research and Innovation) [MRC_UP_1201/10 to E.A.M. and MC_U105185859 to M.M.B.]. For the purpose of open access, the author has applied a CC BY public copyright licence to any Author Accepted Manuscript version arising.

## Funding

This work was supported by funding from the UK Medical Research Council (MRC_UP_1201/10 to E.A.M. and MC_U105185859 to M.M.B.), ALSAC (to M.M.B. and B.S.), Research Grant Council of Hong Kong (No. 26303018 and 16309721 to J.H.), SFPBRNS scheme from the University of Hong Kong (No. 201909185073 to W.L.); and a European Union’s Horizon 2020 research and innovation programme under the Marie Skłodowska-Curie grant agreement (No. 838945 to X.-H.L.).

## Author contributions

X.-H.L. collected sequence data, wrote scripts and performed all the computational analyses, made yeast strains, cloned the plasmids and purified proteins, performed CD, fluorescence, biochemical, imaging, functional, and genetics experiments, C.W.H.Y. and S.M.V.F. performed the NMR experiments and corresponding analyses, N.G.-N. performed biochemistry experiments pertaining localization analyses, V.S. contributed to sample preparations and optimizations of liposome and functional assays, H.Z. performed the Langmuir-monolayer experiments under the supervision of W.L. and J.H., C.G. and B.L. performed experiments related to mitochondrial function under the supervision of M.M.B., M.K. and D.R.G. for helping with experiments and interpretation of mitochondrial functional analysis, A.M. helped with creating yeast strains used in the research, M.W. and K.M. contributed to establishment of the methodology of mitochondrial activity assays, C.M.J. contributed to the interpretation of biophysical characterizations, M.B. aided in computational analysis of disordered protein ensemble, B.S. aided in evolutionary analyses, E.A.M. contributed to design and optimization of yeast genetics experiments. X.-H.L., E.A.M., and M.M.B. designed the research, analyzed and interpreted the results, and wrote the manuscript; X.-H.L., C.W.H.Y., E.A.M., and M.M.B. contributed to data visualization. M.M.B. and X.-H.L. conceived, managed, and set the direction of the project. M.M.B. and E.A.M. co-supervised the project and S.M.V.F. co-supervised the NMR section of the project. All authors contributed to the editing of the manuscript.

## Competing interests

The authors declare no competing interests.

## Data and materials availability

All data are available in the manuscript or the supplementary materials; plasmids and strains described can be obtained from X.-H.L., E.A.M., or M.M.B.

## Notes

### Competing Interest Statement

The authors have declared no competing interest.

