## Extended Data Figures 1-30 for "A molecular mechanism for membrane chaperoning by a late embryogenesis abundant protein"

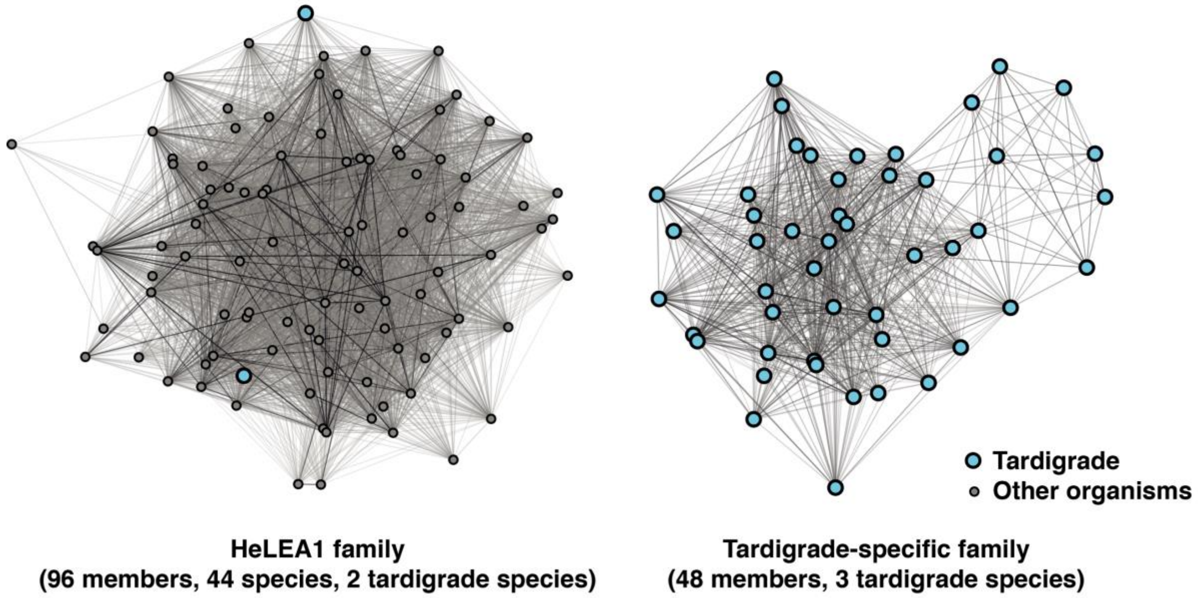


**Extended Data Fig. 1 A comprehensive sequence search identified two homologous protein families for tardigrade IDPs that confer desiccation tolerance when expressed in unicellular organisms.** These include an evolutionarily conserved family (HeLEA1 family, **left**) and a tardigrade specific family (**right**). Homologs from tardigrades are highlighted in blue (**Extended Data Table 1 and Table2**). In this network, nodes represent individual sequences, and the edge weight represents the sequence identity over the aligned region.

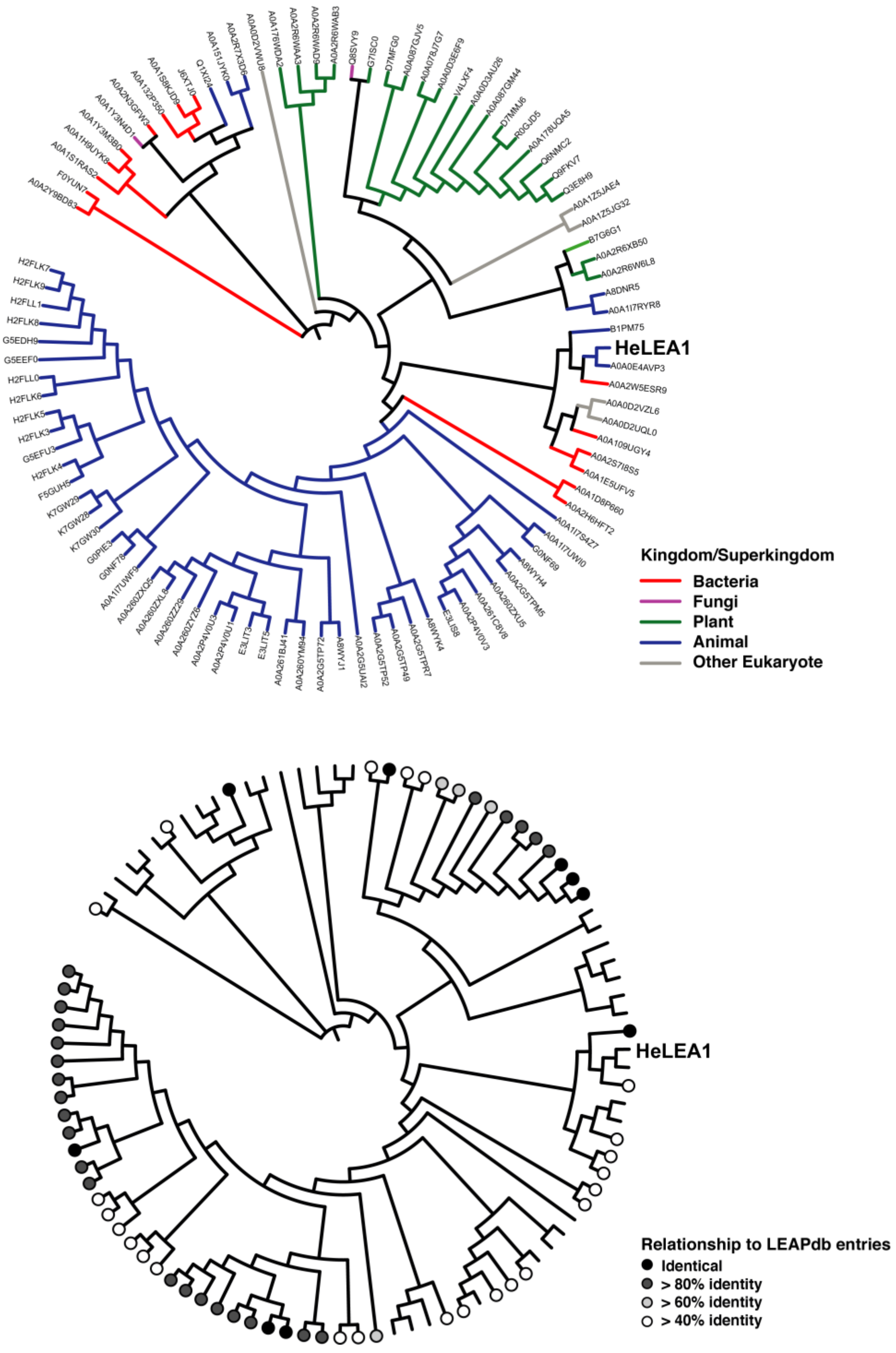


**Extended Data Fig. 2 Evolutionary information for HeLEA1 protein family members annotated by species of origin (top) and sequence similarity to curated entries from the LEA Protein Database (bottom).** In the top panel, each homolog is represented by its UniProt ID. The HeLEA1 homologs belong to 44 different species across five kingdoms or superkingdoms, illustrating the conservation of sequence features over millions of years of evolution. The evolutionary tree is identical to that shown in Fig. 1a. Of the 96 identified homologs of HeLEA1, 59 have regions that share over 40% identity to the protein entries in the LEA Protein Database.

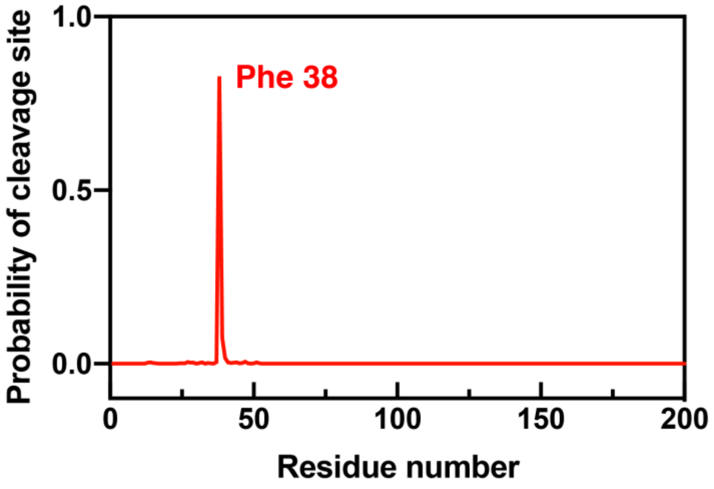


**Extended Data Fig. 3 Prediction of location of the cleavage site of mitochondrial-targeting sequence by TargetP.** This suggests that full-length HeLEA1 has a high probability of being cleaved at phenylalanine 38 by the mitochondrial processing peptidase inside mitochondrial matrix.

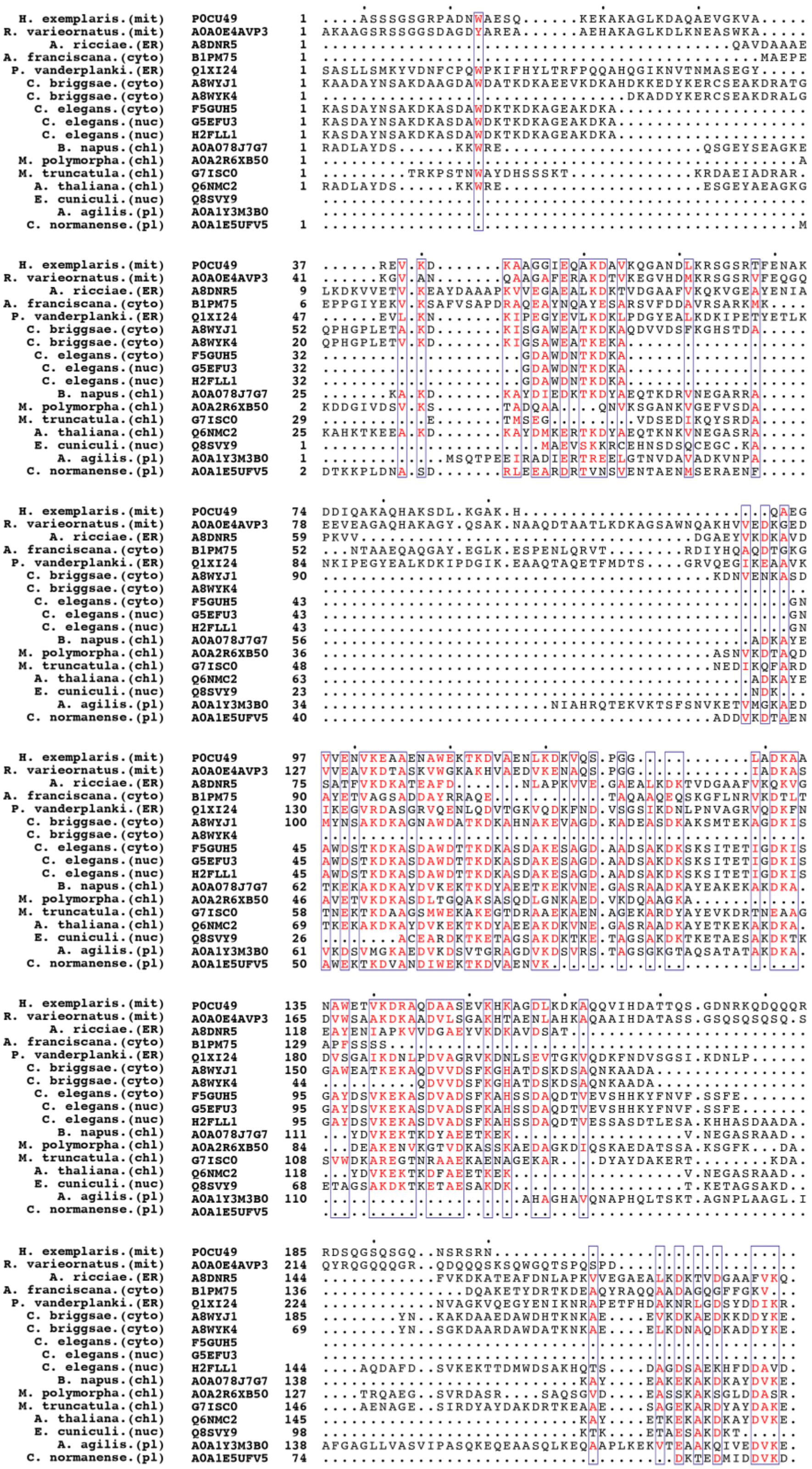

**Extended Data Fig. 4 Sequence alignment of HeLEA1 and its representative homologs after removing predicted signal sequences.** For each sequence, the species of origin, predicted localization, and UniProt ID are indicated. Alignment is visualized using ESPript, with %Equivalent coloring scheme, positions with global equivalent score over 0.5 are highlighted.

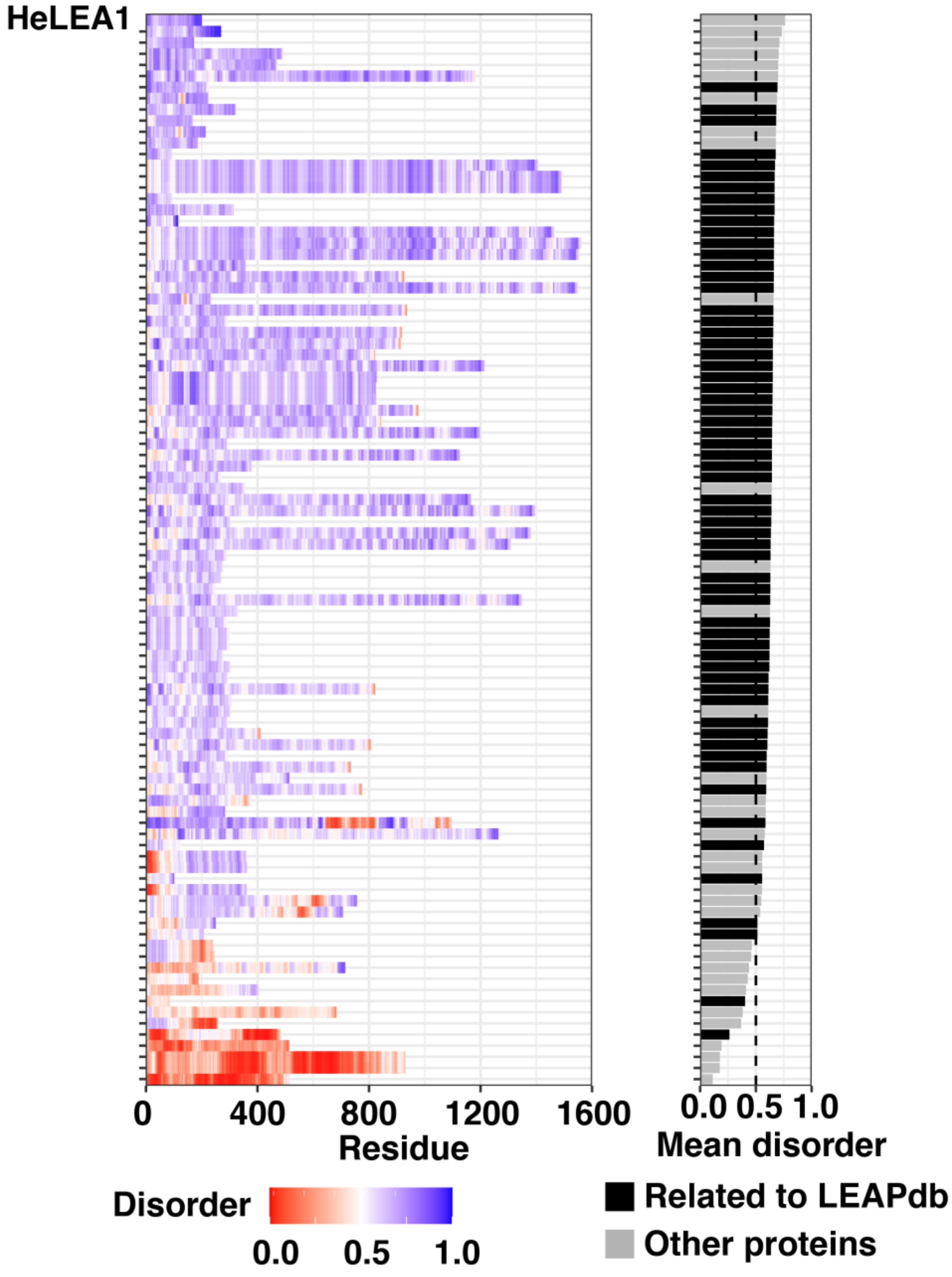


**Extended Data Fig. 5 Residue-specific disorder prediction for 96 HeLEA1 homologs using IUPred2A (left) sorted by the mean disorder scores across their sequences (right).** Homologs related to the recorded entries in the LEA protein database (LEAPdb) are highlighted in black. 83 of 96 with a mean IUPred disorder score > 0.5. HeLEA1 have the highest mean disorder propensity.

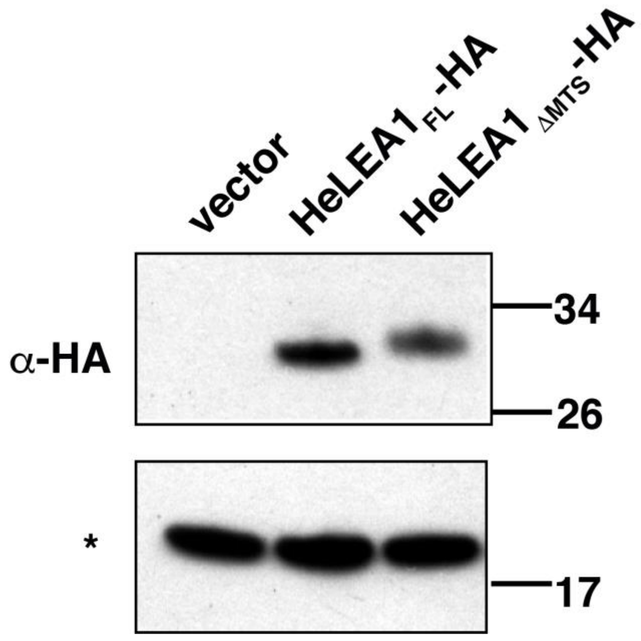


**Extended Data Fig. 6 Immunoblot of different HeLEA1 constructs expressed in yeast at steady phase.** Cells expressing HA-tagged HeLEA1$_{FL}$ migrated as one band, which is slightly lower than HeLEA1$_{\Delta MTS}$-HA migrated.

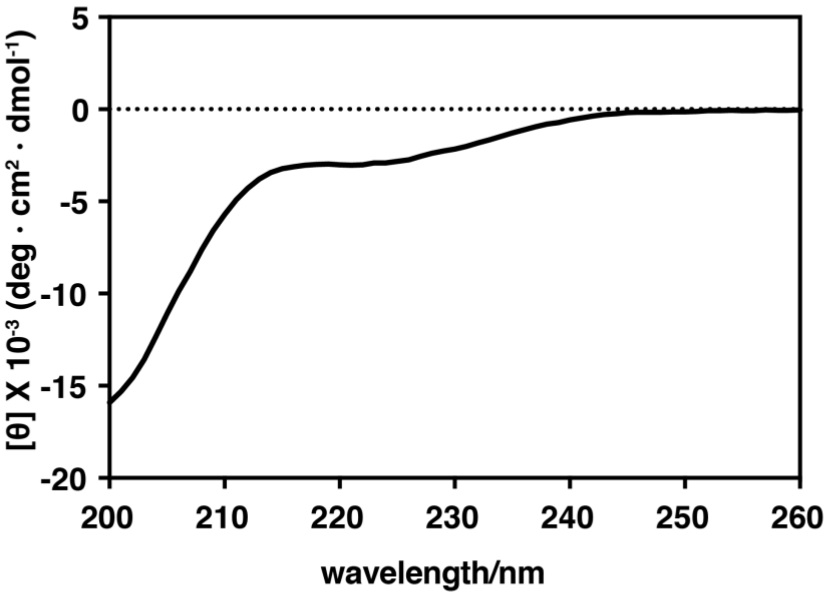


**Extended Data Fig. 7 Circular dichroism spectroscopy of HeLEA1 in solution.** The spectrum lacks the characteristic signals of alpha-helical (negative peaks at 222 nm and 208 nm) and beta-sheet (negative peak at 218 nm) secondary structures and has a characteristic signal of a disordered protein (negative peak at 200 nm).

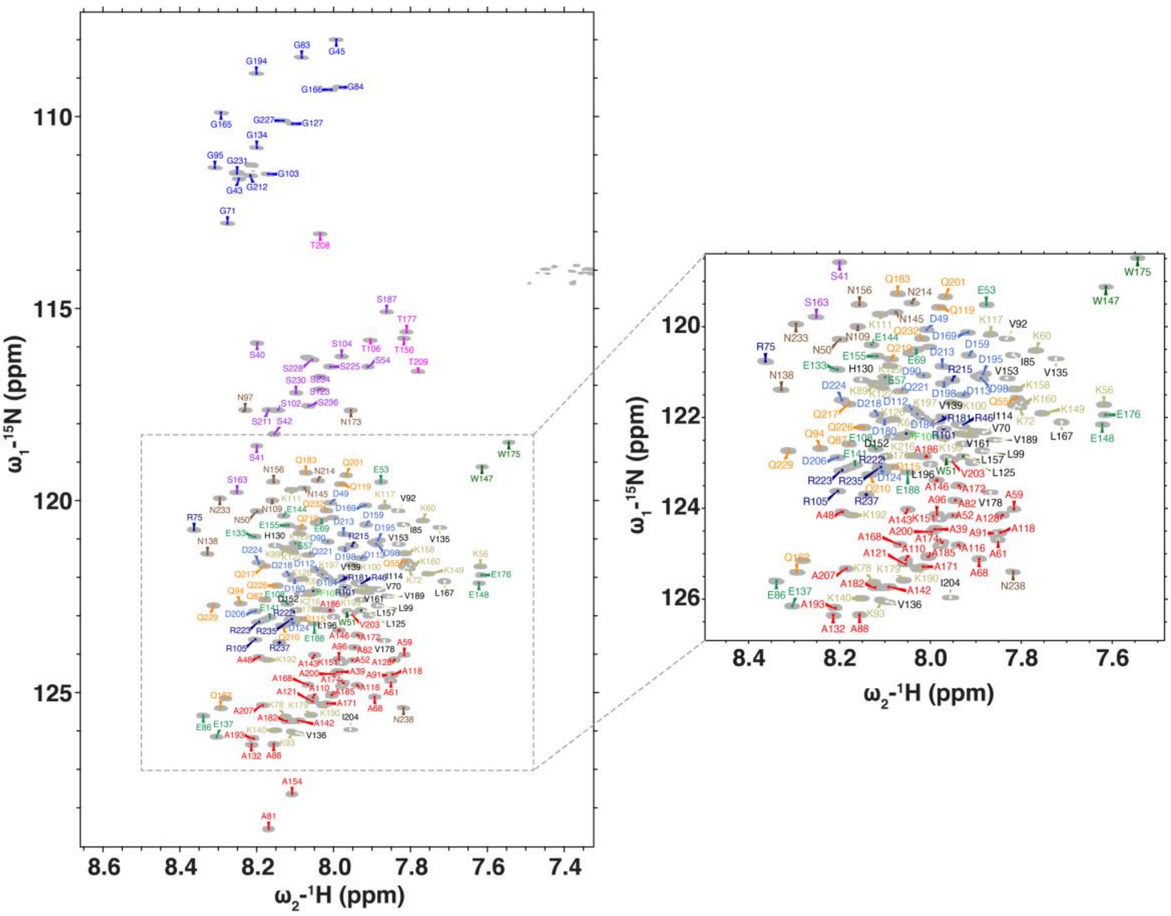


**Extended Data Fig. 8 Full backbone assignment of $^{1}H_{N}$,$^{15}N$ 2D HSQC of HeLEA1.** The resonances are coloured based on their corresponding amino acid type and numbered according to the numbering in full-length HeLEA1.

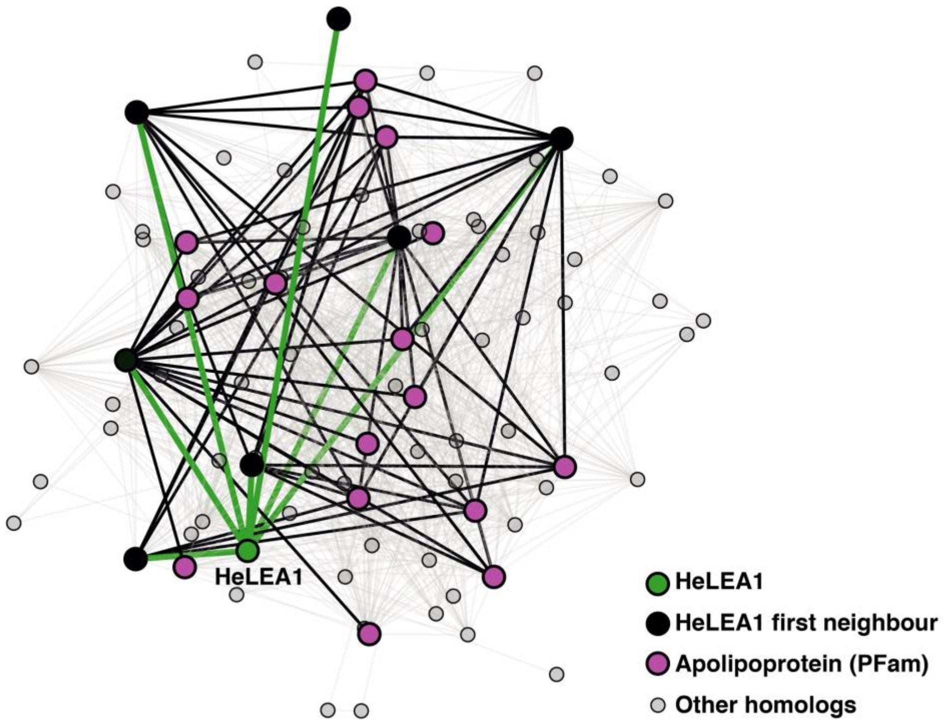


**Extended Data Fig. 9 HeLEA1 shares distant homology with annotated apolipoproteins.** Network constructed by pairwise sequence similarity was further filtered for pairs of homologs with > 40% identity. HeLEA1 (green) shares low similarity with an annotated apolipoprotein (magenta). However, six of the seven first neighbors of HeLEA1 (black) share a reasonable level of homology with apolipoproteins annotated in Pfam. The edges, which represent > 40% pairwise sequence identity in aligned regions, are highlighted in green (between HeLEA1 and its first neighbors) or black (between HeLEA1 first neighbors and annotated apolipoproteins).

```
       1        10        20        30             40        50
HeLEA1 ASSSGSGRPADNWAESQKEKAKAGLKDAQAEVGKVA.....REVKDKAAGGIEQA
1EQ1   ...............EMEKHAKEFQKTFSEQFNSLVNSKNTQDFNKALKDGSDSV

                60        70        80        90       100
HeLEA1 KDAVKQGANDLKRSGSRTFENAKDDIQAKAQHAKSDLKGAKHQAEGVVENVKEAA
1EQ1   LQQLSAFSSSLQGAISDANGKAKEALEQARQNVEKTAEELRKAHPDVEKEANAFK

           110       120       130       140       150       160
HeLEA1 ENAWEKTKDVAENLKDKVQSPGGLADKAANAWETVKDRAQDAASEVKHKAGDLKD
1EQ1   DKLQAAVQTTVQESQKL.......AKEVASNMEETNKKLAPKIKQAYDDFVKHAE

               170       180       190       200
HeLEA1 KAQQVIHDATTQSGDNRKQDQQQRRDSQGSQSGQNSRSRN
1EQ1   EVQKKLHEA...............................
```

| PDB ID | Coverage | Confidence | Identity | Pfam annotation |
|---|---|---|---|---|
| 1EQ1 | 16-169 (74%) | 81.4% | 10% | PF07464: Apolipophorin-III |
| 2X43 | 45-94 (24%) | 78.2% | 10% | Unclassified |
| 1XQ8 | 8-98 (44%) | 50.5% | 11% | PF01387: Synuclein |
| 2L9Q | 91-177 (43%) | 43.0% | 20% | PF04119: Heat shock protein 9/12 |

**Extended Data Fig. 10 HeLEA1 has weak sequence homology with lipid-interacting proteins. Top:** Sequence alignment between mature HeLEA1 and the top hit from a PhyRE2 homology search (PDB: 1EQ1), illustrating the very weak sequence similarity but good coverage of the sequence. **Bottom:** Summary of hits from the PhyRE2 search. Three of four hits (1EQ1, 1XQ8 and 2L9Q) contain the Pfam annotation of protein families involved in lipid interaction, although with low confidence to identify them as proper structural scaffolds for HeLEA1 (< 90%).

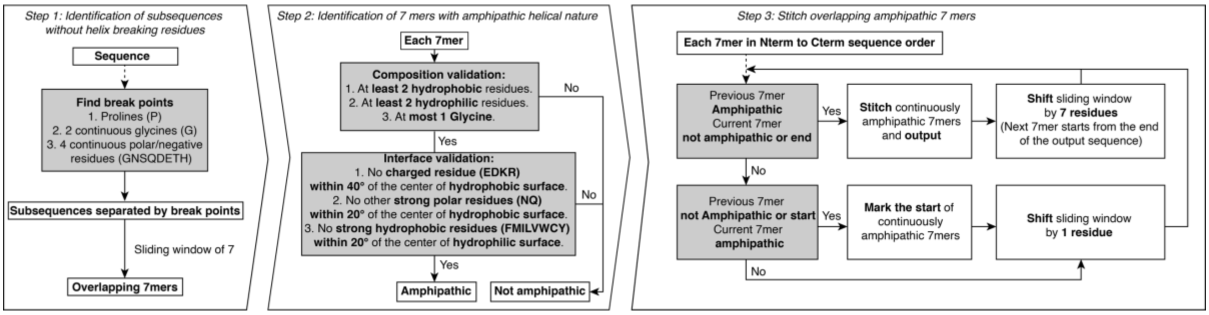


**Extended Data Fig. 11 Bioinformatics pipeline for discovery of amphipathic 3-11 helical motifs in a given protein, with a minimal length of 7.**

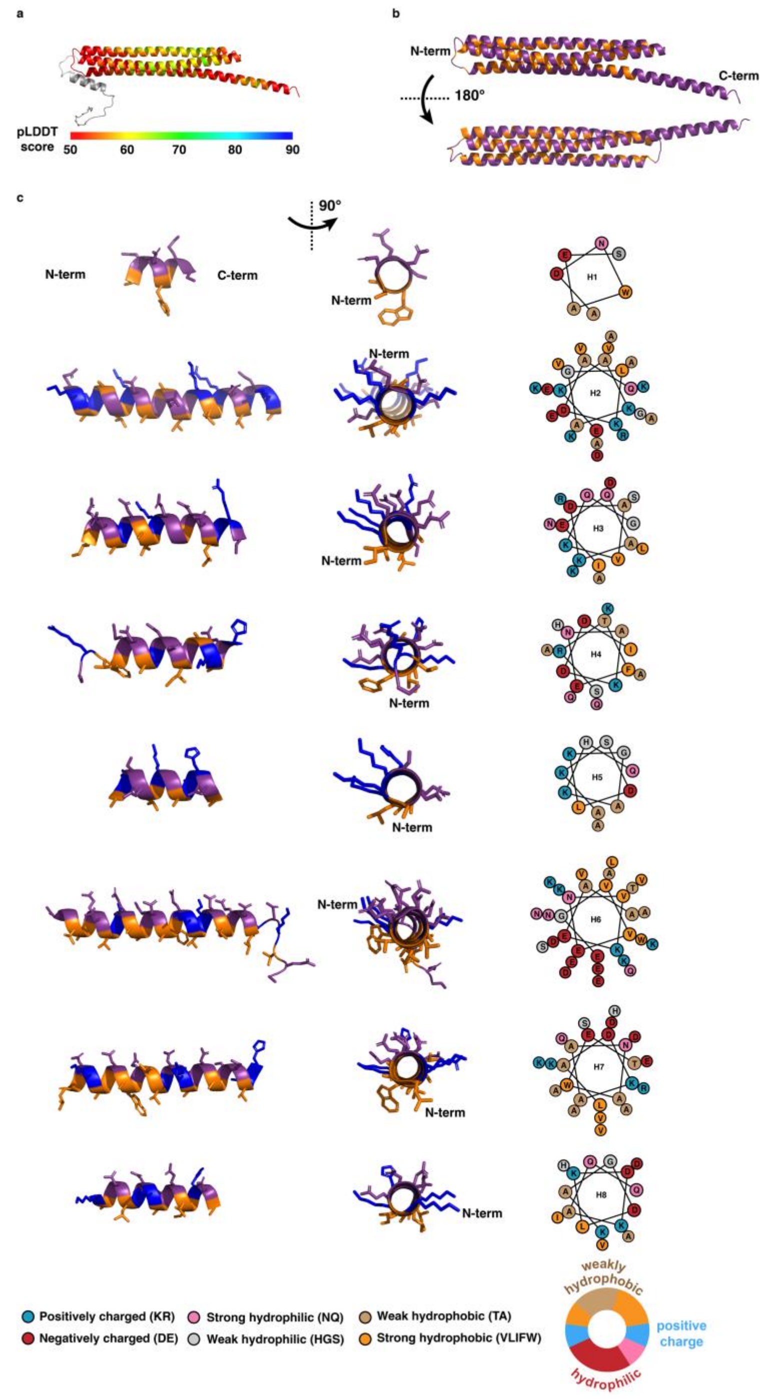


**Extended Data Fig. 12 Predicted structure of HeLEA1. a,** AlphaFold2 prediction of $HeLEA1_{FL}$, rendered by pLDDT score. The N-terminal MTS is rendered in gray. The low confidence score suggests the global higher-order structure is less reliable. **b,** AlphaFold2 prediction of HeLEA1, rendered by residue hydrophobicity. Hydrophobic residues (Ala, Cys, Phe, Ile, Leu, Met, Pro, Thr, Val, Trp, Tyr), are colored in orange, the rest of residues in purple. It is clear the AlphaFold predicted tertiary structure exhibits mismatching of amphipathic surfaces, and the tertiary structure is probably biased by the absence of membrane causing attraction between hydrophobic surfaces. **c,** Amphipathic elements predicted by biophysical criteria display coherent amphipathic surfaces.

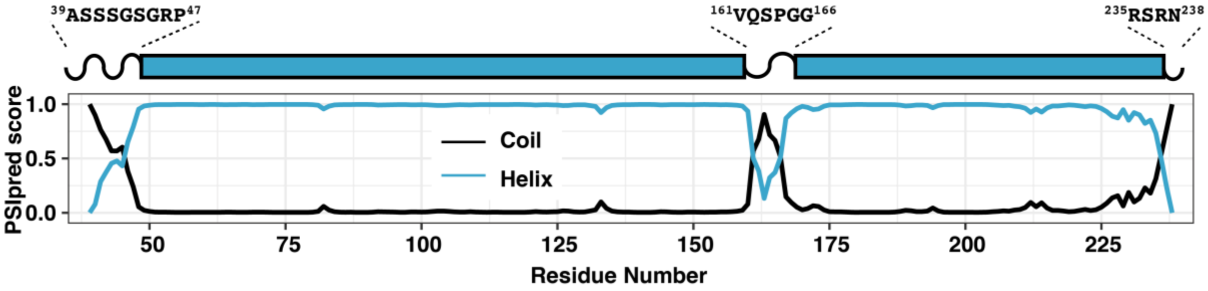


**Extended Data Fig. 13 PSIPRED data for secondary structure propensity in its functional form of HeLEA1.** Most of the residues in HeLEA1 are predicted to be helical with high confidence.

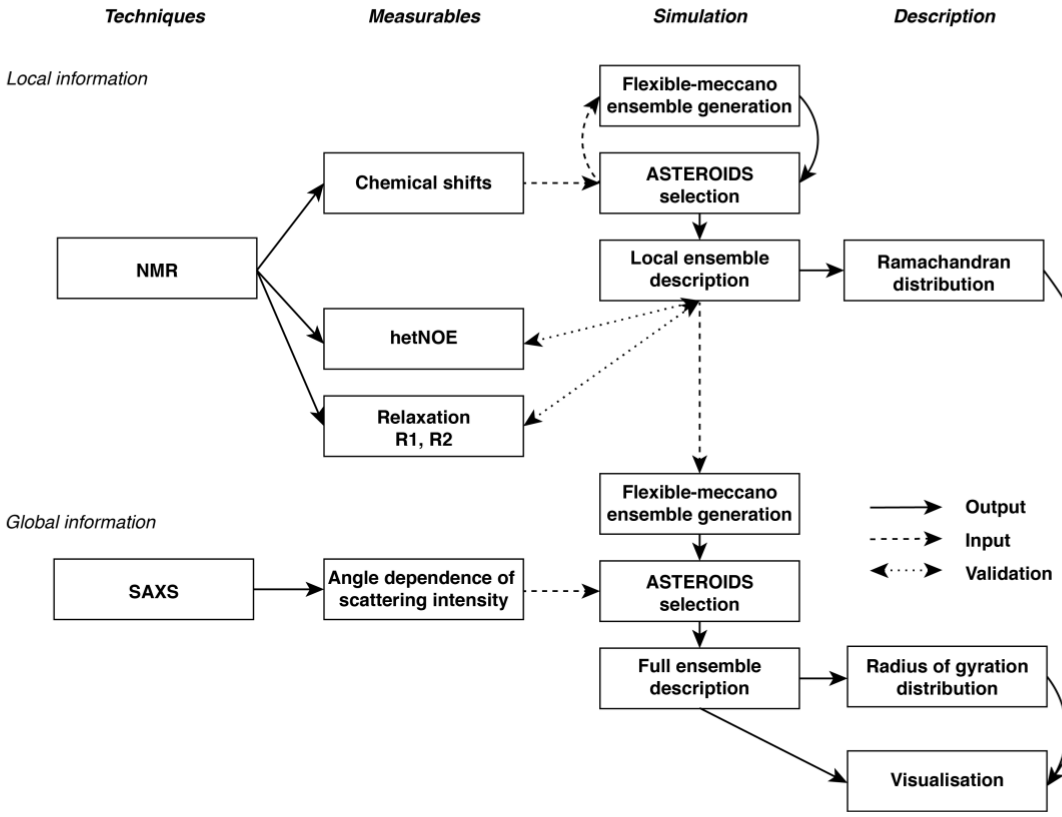


**Extended Data Fig. 14 Elaborated integrative simulation pipeline for the solution state of HeLEA1.**

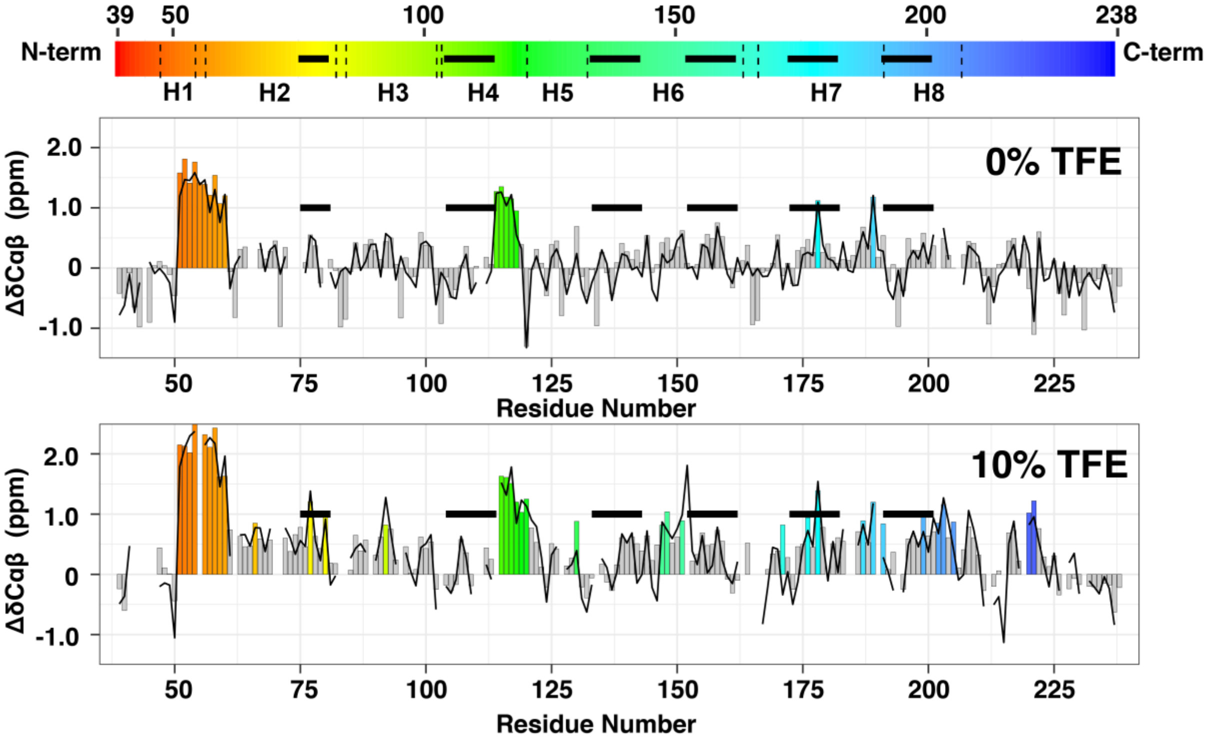


**Extended Data Fig. 15 Secondary chemical shift ($\Delta\delta C_{\alpha\beta}$) of HeLEA1 in 0% TFE (top) or 10% TFE (bottom).** Black line represents *ASTEROIDS* fit of $\Delta\delta C_{\alpha\beta}$. Residues with significant positive secondary chemical shifts (> 0.8 ppm) are highlighted by color. Black bars represent conserved LEA motifs mapped in **Fig. 1**.

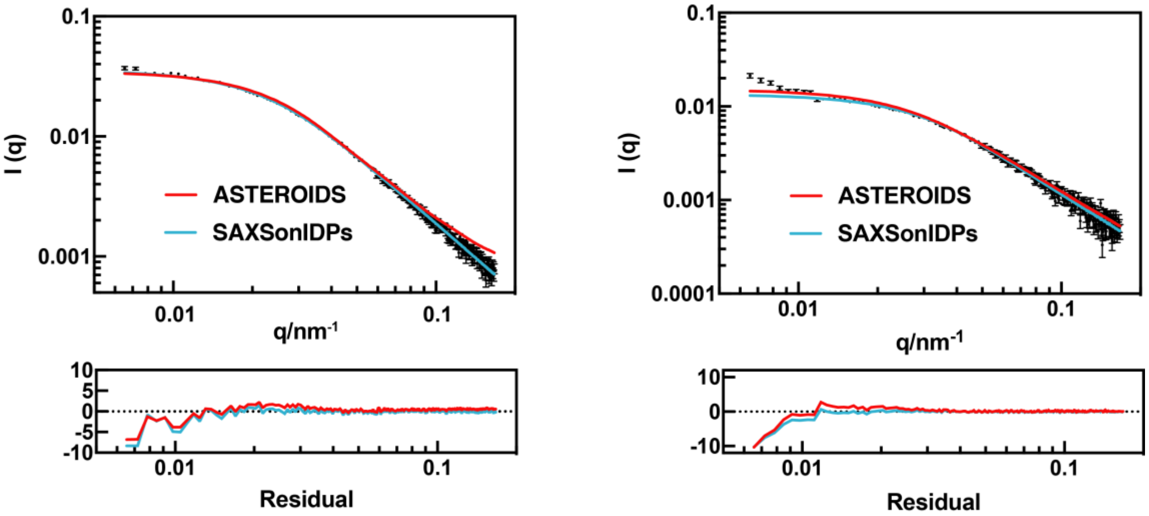


**Extended Data Fig. 16 SAXS data and fitting of HeLEA1 in 0% TFE (left) and 10% TFE (right) using two different fitting methods.** The ensemble estimation by ASTEROIDS agreed well with the *SAXSonIDPs*[34] fitting. The residuals are standardized. SAXS data are represented as the mean ± SE (**Extended Data Table 4**).

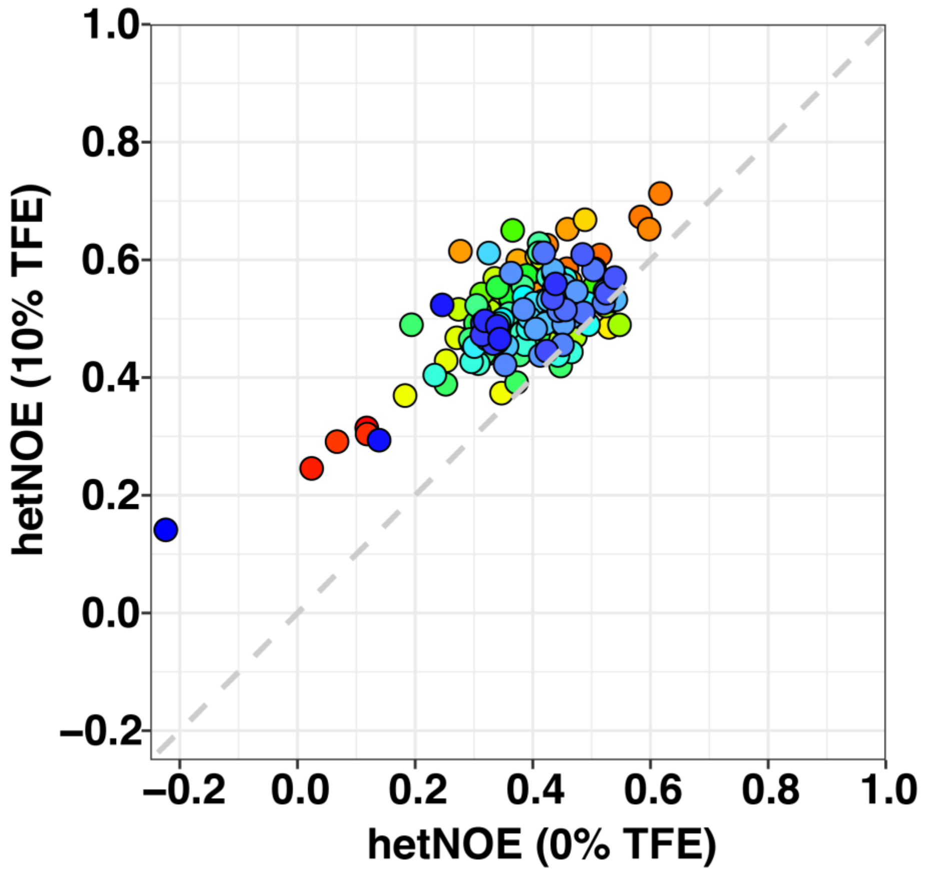


**Extended Data Fig. 17 Correlation of residue-specific $^{15}N\{^{1}H\}$ heteronuclear NOE (hetNOE) for HeLEA1 in 0% or 10% TFE.** A global increase of hetNOE were observed as a result of 10% TFE. The coloring scheme is the same as in **Fig. 2b** and **Extended Data Fig. 15**.

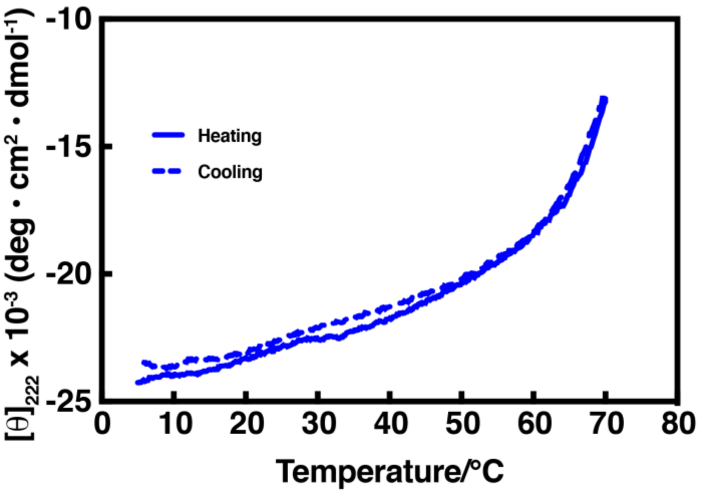


**Extended Data Fig. 18** Thermal melting and refolding curves monitoring dynamics of helicity of HeLEA1 induced by 1mM POPS SUVs. The helical state is in fast equilibrium with the disordered state and the folding process is completely reversible, corresponding well to a dynamic and environmental-sensitive disorder-to-helical transition process.

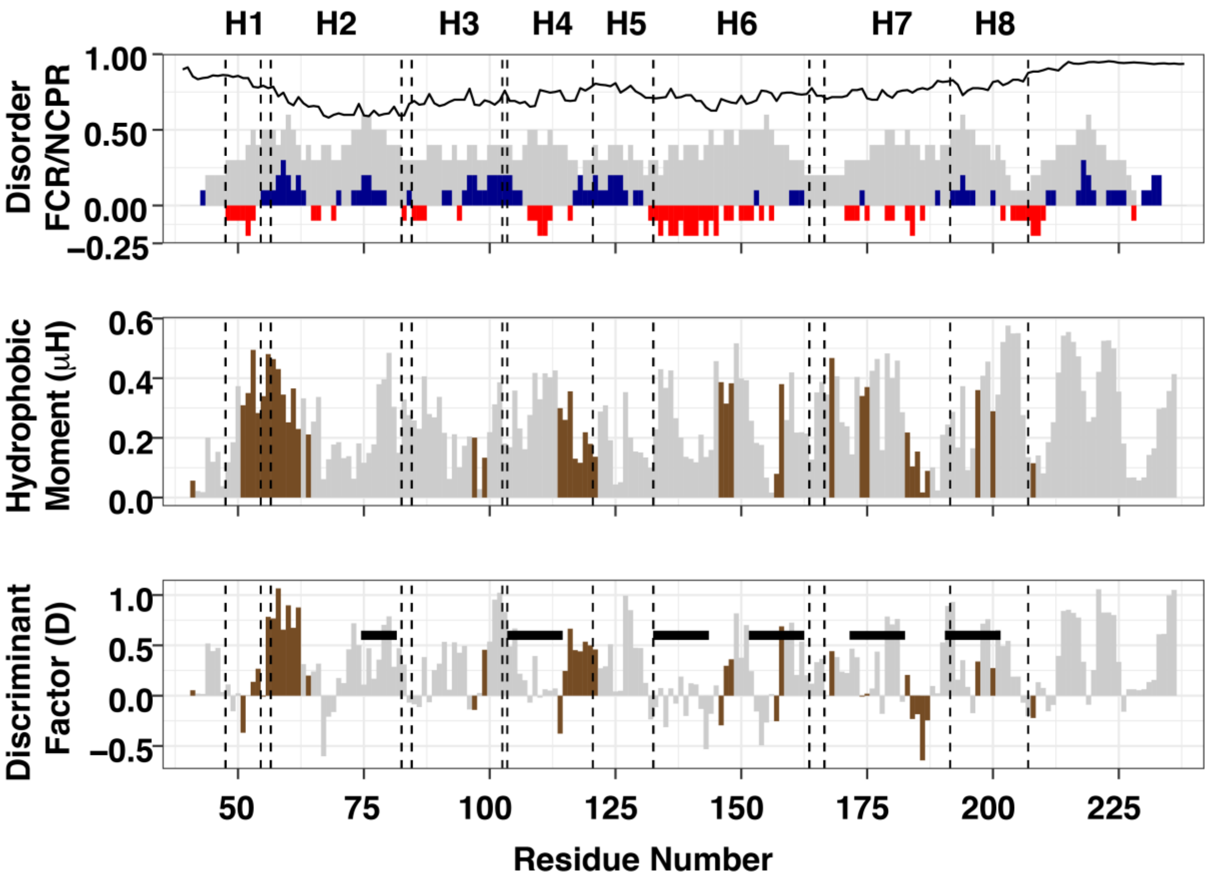


**Extended Data Fig. 19 Sequence properties of HeLEA1. Top**, plot showing IUPred2 predicted disorder propensity (black line), fraction of charged residues (FCR, gray bars), and net charge per residue (NCPR, red/blue bars for net negative/positive charges). **Middle**, distribution of hydrophobic moments of local short amphipathic elements (length of 5). **Bottom**, distribution of lipid binding discriminant factor D of local short amphipathic elements (length of 5), which integrates contribution from hydrophobic moments and charge interactions; black bars indicate conserved LEA motifs mapped in **Fig. 1**. Regions with increased helical propensity in 10% TFE corresponding to colored residues in **Fig. 2b** are highlighted in brown for hydrophobic moment and lipid binding factor plots.

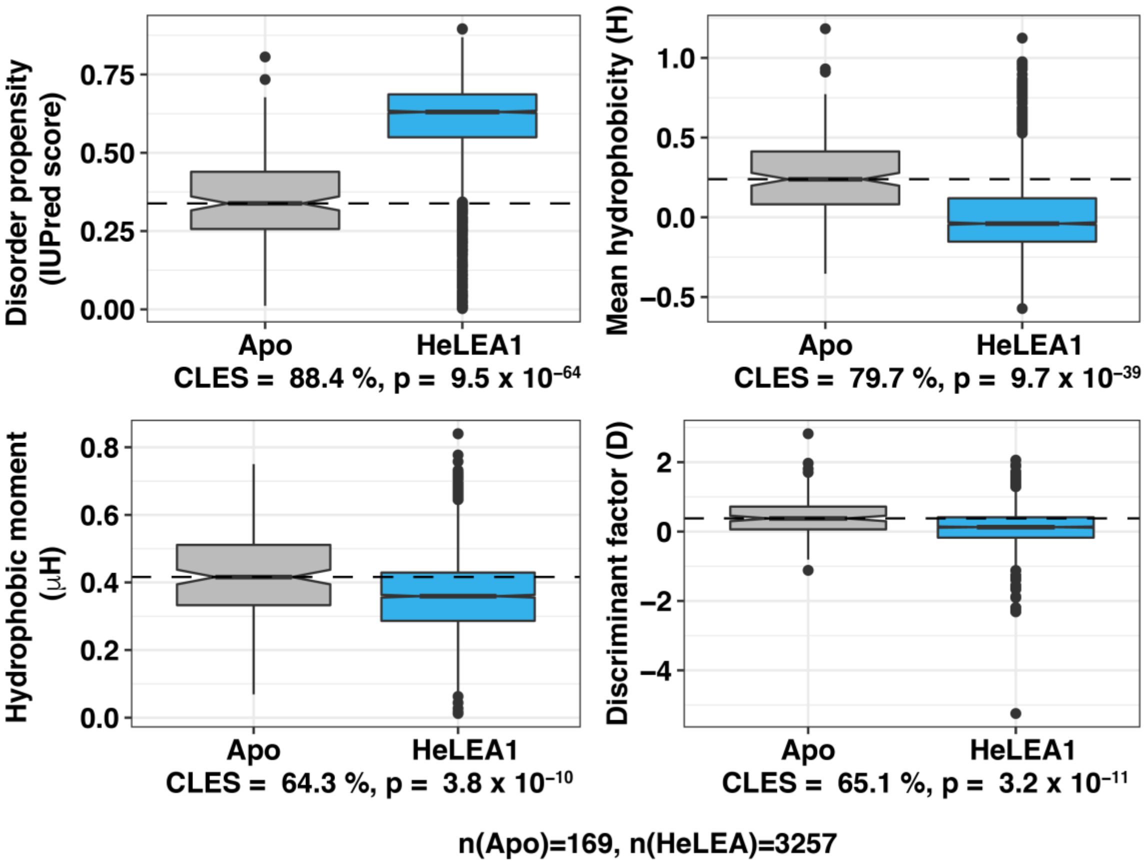


**Extended Data Fig. 20** HeLEA1 homologs have distinct features that discriminate them from classic lipid-binding helices in apolipoproteins. Amphipathic 3-11 helices in HeLEA1 homologs possess a significantly higher propensity of being intrinsically disordered, less hydrophobic (lower mean hydrophobicity) and amphipathic (lower hydrophobic moment), and has a lower discriminant score D, indicating weaker lipid-binding ability. Each box plot demonstrates specific biophysical parameters of 3-11 amphipathic helices either from human apolipoprotein family (Apo) or HeLEA1 homologs (HeLEA1), using the algorithm described in **Extended Data Fig. 11**. Each box represents the IQR of the dataset, whiskers represent plus/minus 1.5 IQR from the box hinge, and outliers are plotted as dot.

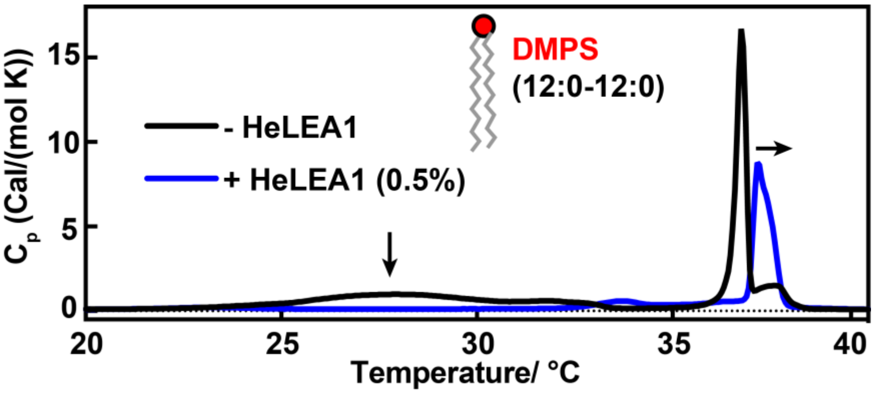


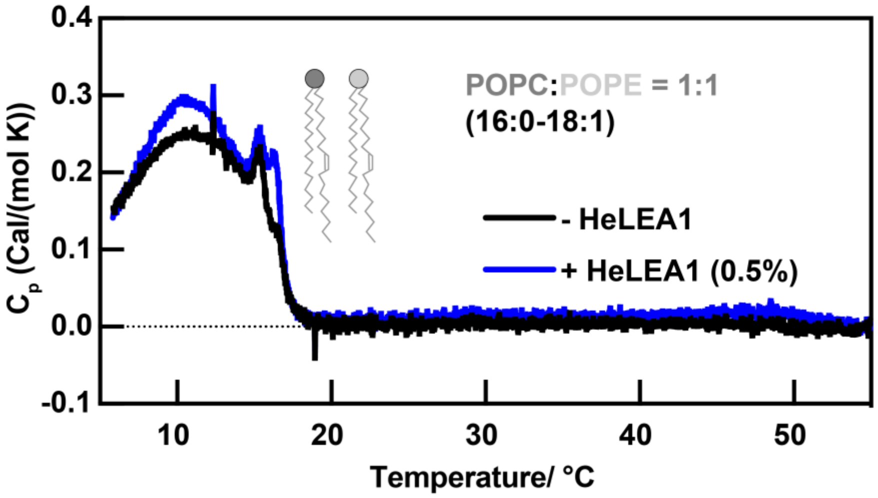


**Extended Data Fig. 21 Differential scanning calorimetry for different lipid compositions. Top,** saturated DMPS SUVs without (black line) and with mature HeLEA1 (0.5% molar ratio, blue line). The addition of HeLEA1 suppressed thermally induced phase transition at low temperature and (broad peaks indicated by arrow) increased phase transition temperature (peak position). **Bottom,** SUVs made with a mixture of neutral lipids POPC and POPE (1:1 molar ratio) without (black line) and with HeLEA1 (molar ratio 0.5%, blue line). Despite minor changes, the addition of HeLEA1 did not induce significant changes in the phase behavior of the POPC/POPE SUVs and the shape of the DSC curve remains largely unchanged. This result corresponds well with very weak interaction between HeLEA1 and the neutral vesicles.

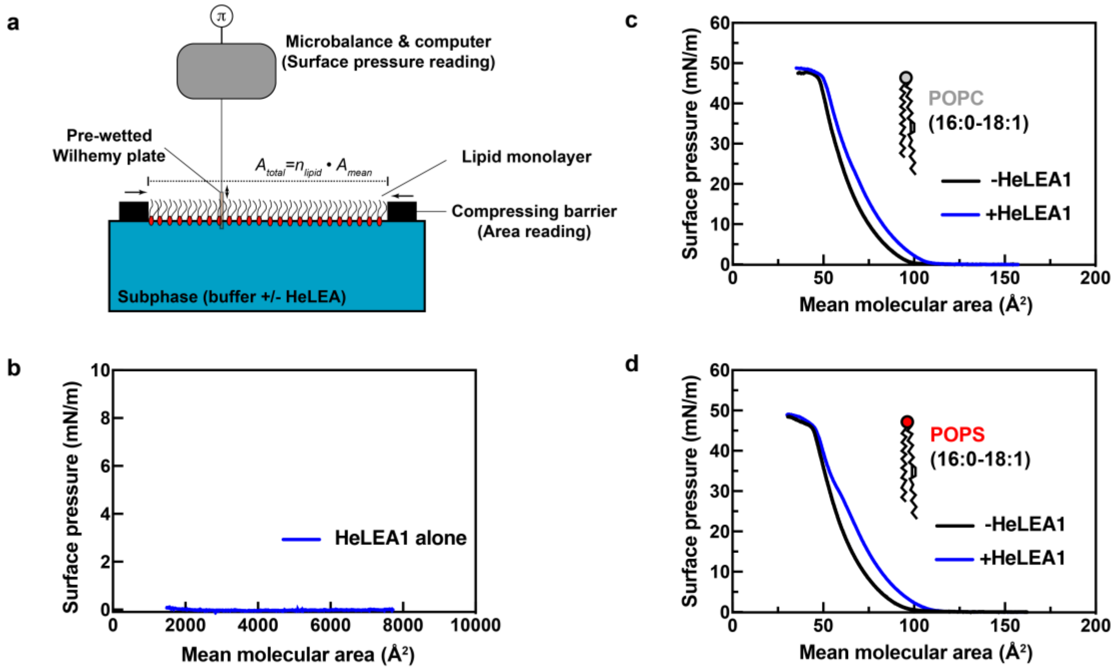


**Extended Data Fig. 22 Langmuir-monolayer methodology characterizing the surface behavior of lipids at an air–water interface. a**, Scheme illustrating the instrument setup for measuring lipid monolayer compression isotherms. Mean molecular area $A_{mean}$ and surface pressure □ are plotted. The subsequently calculated $C_S{}^{-1}$ values are reported in **Fig. 3e**. **b-d,** Compression isotherm of **b,** HeLEA1 alone, **c,** POPC monolayer with or without HeLEA1 and **d,** POPS with or without HeLEA1. The plots revealed that 3 nM HeLEA1 alone induces a negligible change in the surface pressure at the air–buffer interface but changes the isotherms of lipid monolayers, suggesting that the observed change in isotherms in POPC and POPS monolayers is not due to the surface effect of HeLEA1 alone.

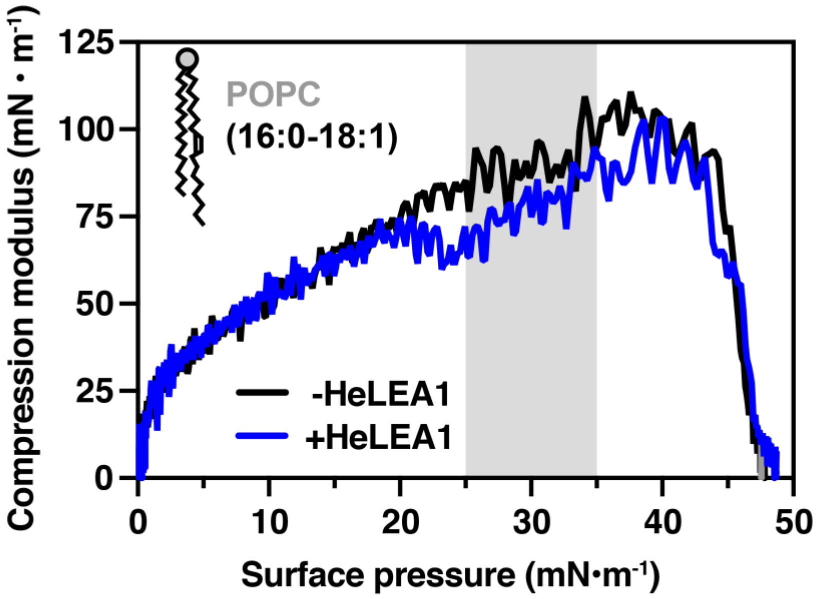


**Extended Data Fig. 23** Change of ${C_S}^{-1}$ values in neutral POPC monolayers with respect to surface pressure, as measured by Langmuir-monolayer methodology, with or without 3nM HeLEA1 (1:50 protein:lipid ratio). Gray areas depict the range of physiological surface pressure.

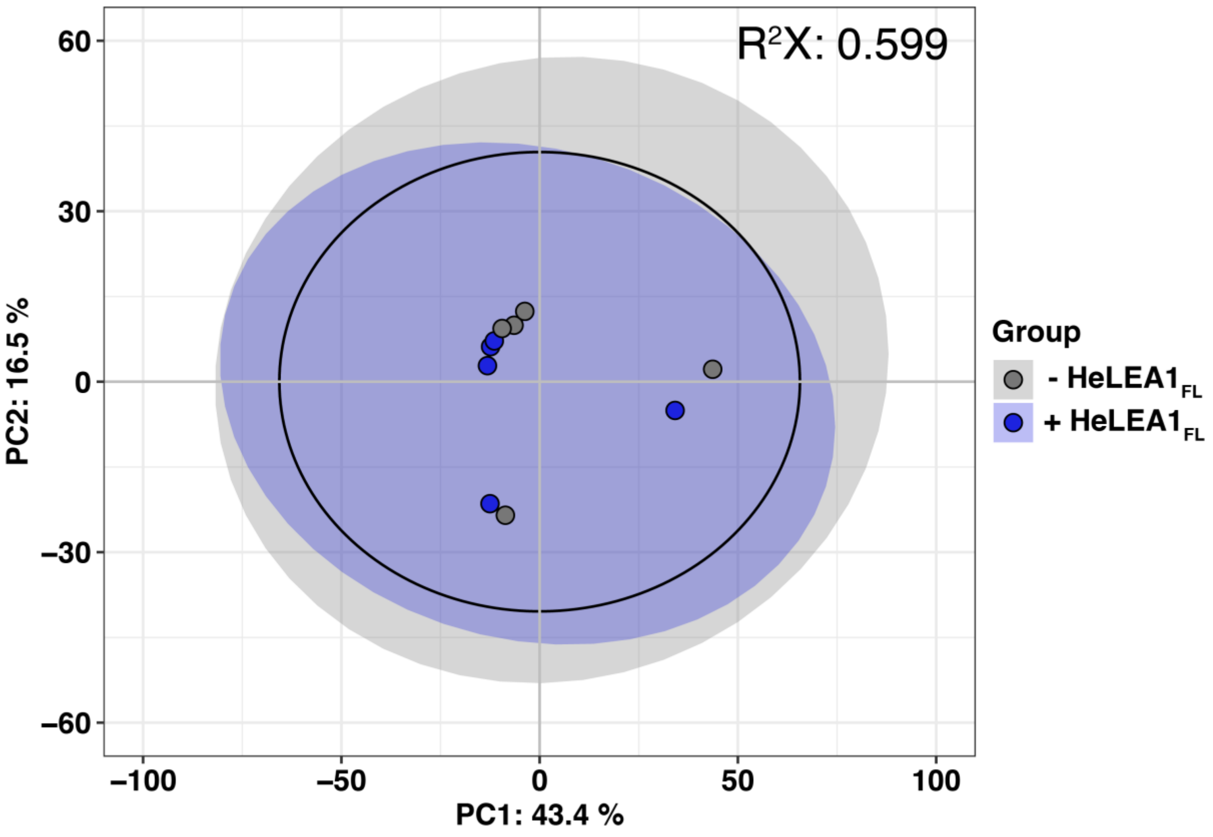


**Extended Data Fig. 24** Principle component analysis of lipidomics data from mitochondria purified from yeast cells with (blue) or without (gray) $HeLEA1_{FL}$ expression grown in normal YPD media. The shaded ellipses represent distribution of each group with a confidence interval of 95%. The black circle represents 95% confidence interval of Hotelling's $T^2$ statistics. The PCA analysis suggests HeLEA1 does not trigger significant change in lipid composition.

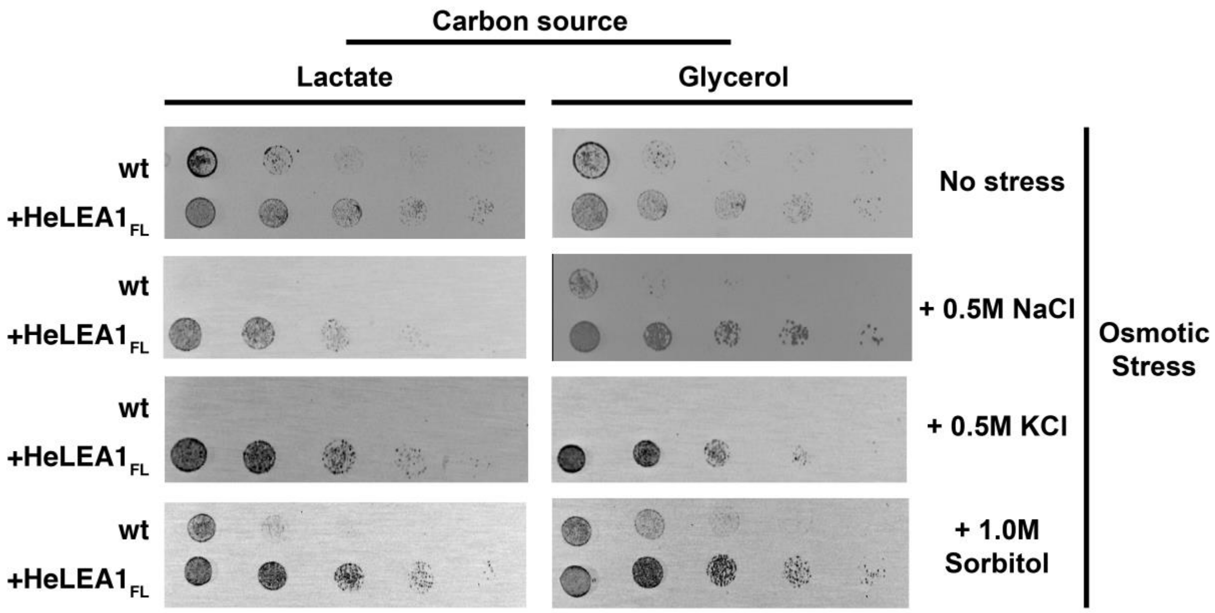


**Extended Data Fig. 25** HeLEA1$_{FL}$ expression confers tolerance to multiple osmotic stresses on non-fermentable carbon sources. Lactate and glycerol which are both non-fermentable carbon sources were combined with 0.5M NaCl, 0.5M KCl, or 1.0M Sorbitol to mimic varios osmotic stresses. In either case, expression of HeLEA1$_{FL}$ confers a significant growth advantage compared to strains without HeLEA1$_{FL}$ expression at 37 °C.

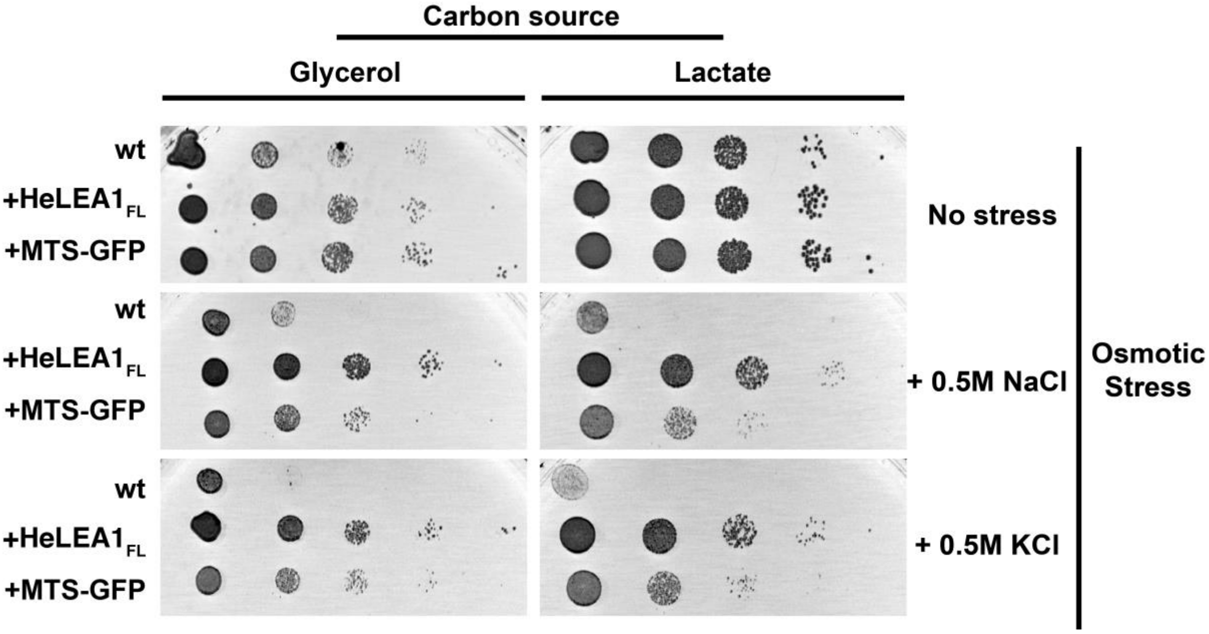


**Extended Data Fig. 26** Hyperosmotic stress tolerance conferred by HeLEA1 is not due to mitochondrial import stress response. Cells overexpressing HeLEA1$_{FL}$ or MTS-GFP were spotted under different osmotic stress on non-fermentable carbon sources at 37 °C. In either case, overexpression and import of MTS-GFP conferred some level of stress tolerance, but is less significant than HeLEA1$_{FL}$.

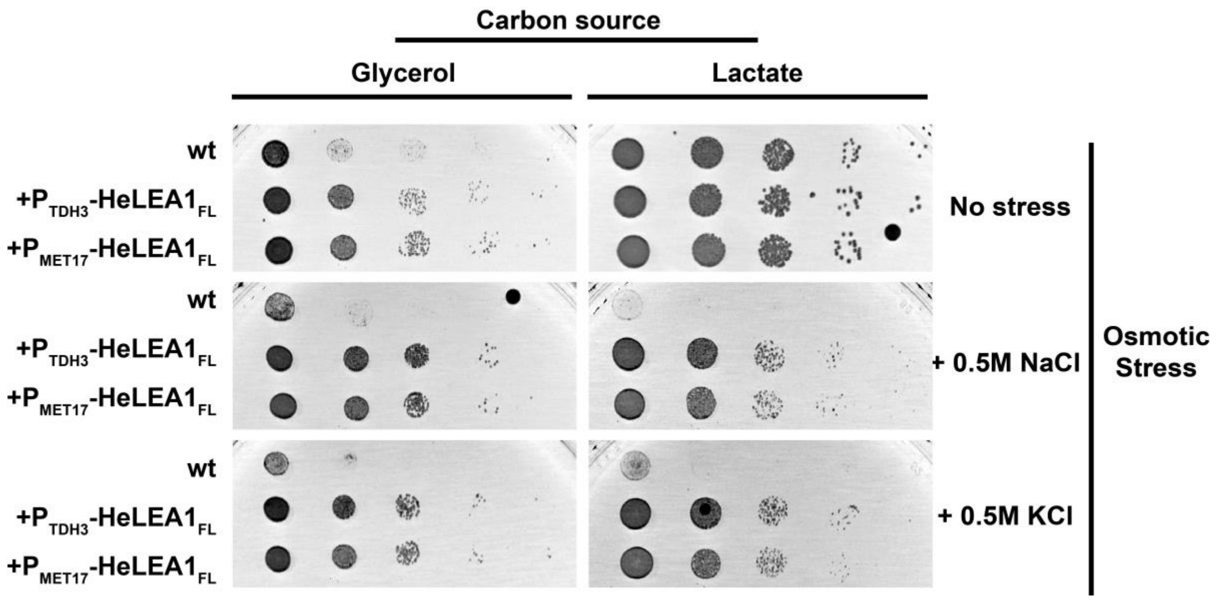


**Extended Data Fig. 27** Hyperosmotic stress tolerance conferred by HeLEA1 is not due to overexpression stress. Cells expressing HeLEA1$_{FL}$ under either a constitutive strong promoter TDH3 or leakage expression from weak promoter MET17 were spotted under different osmotic stress on non-fermentable carbon sources at 37 °C. In either case, expression of HeLEA1$_{FL}$ conferred similar stress tolerance phenotype.

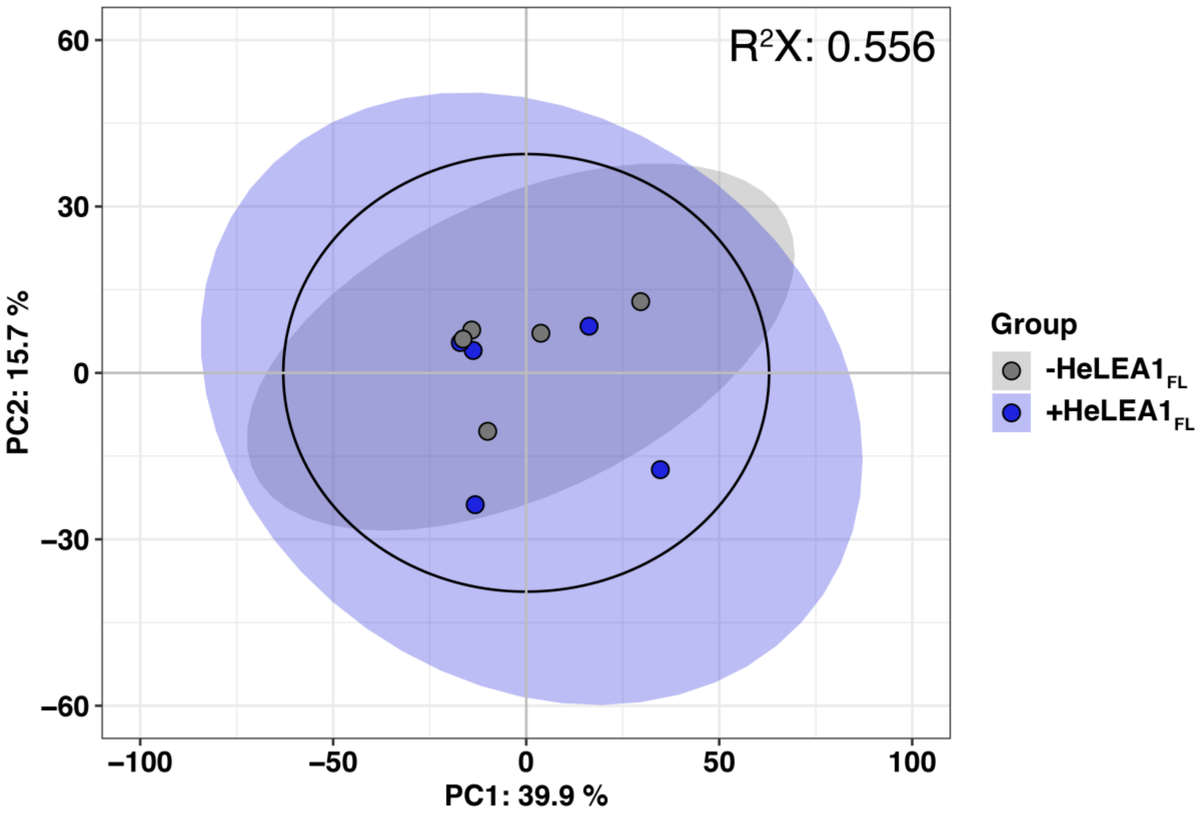


**Extended Data Fig. 28** Principle component analysis of lipidomics data from mitochondria purified from yeast cells with (blue) or without (gray) HeLEA1$_{FL}$ expression grown in non-fermentable YEP+Glycerol media (3%). The shaded ellipses represent distribution of each group with a confidence interval of 95%. The black circle represents 95% confidence interval of Hotelling's $T^2$ statistics. The PCA analysis suggests HeLEA1$_{FL}$ does not trigger significant change in lipid composition under non-fermentable condition.

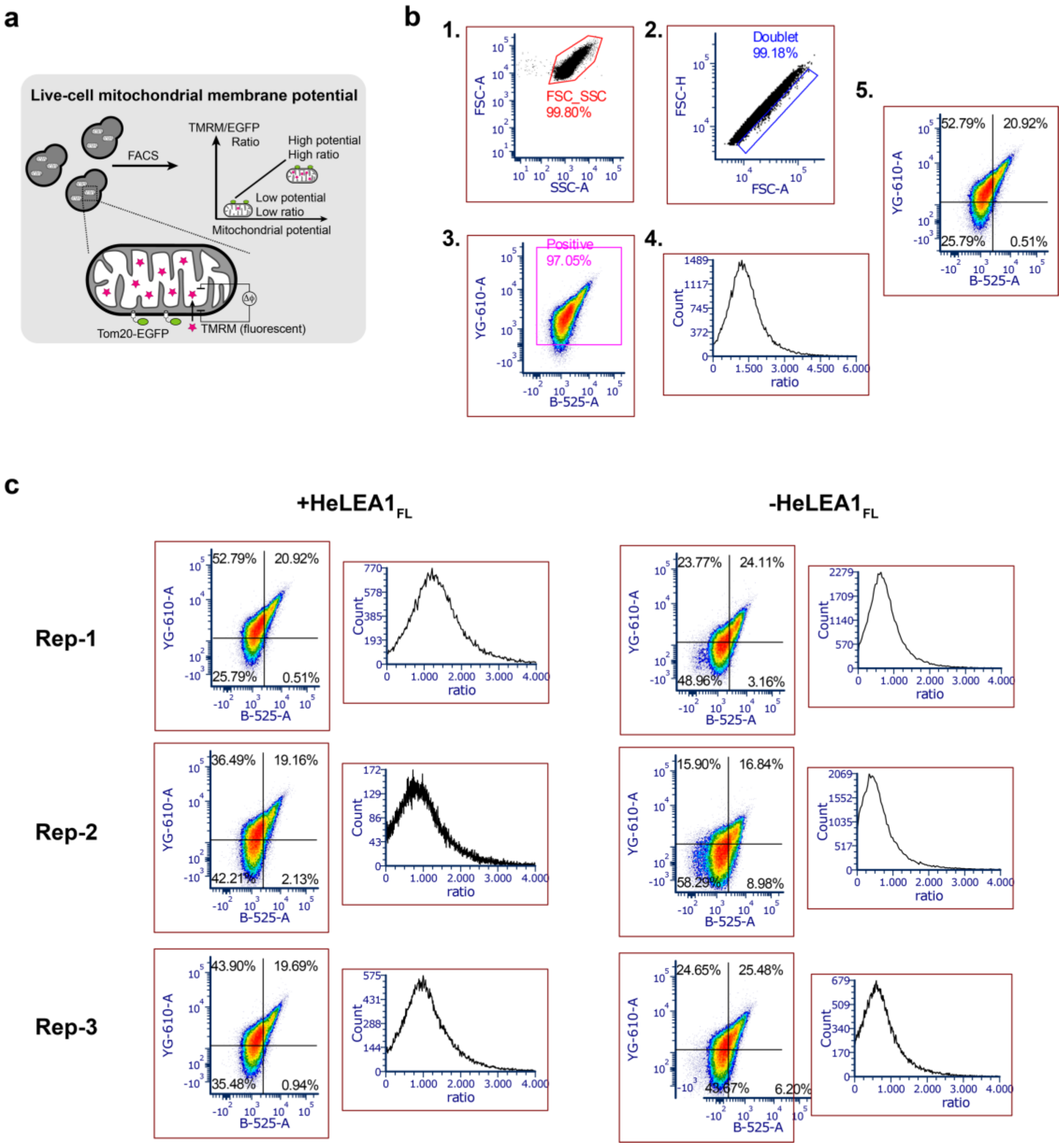


**Extended Data Fig. 29 HeLEA1$_{FL}$ expression enhances mitochondrial membrane potential on fermentable carbon source. a,** Scheme illustrating the principle of TMRM-based assessment of mitochondrial membrane potential. **b,** Gating strategy in the assay first gates away dead cells by a polygon gate on front scattering-side scattering plot (1), followed by doublet exclusion on front scattering height-area (2). The data is either visualized by quadrant percentage (5), or further represented as histogram (4) after gating on positive TMRM and GFP signal (3). **c,** FACS data of three independent replicates for cells with or without HeLEA1$_{FL}$ expression. The summary statistics for ratio between TMRM and GFP signal is represented in **Fig. 4d**.

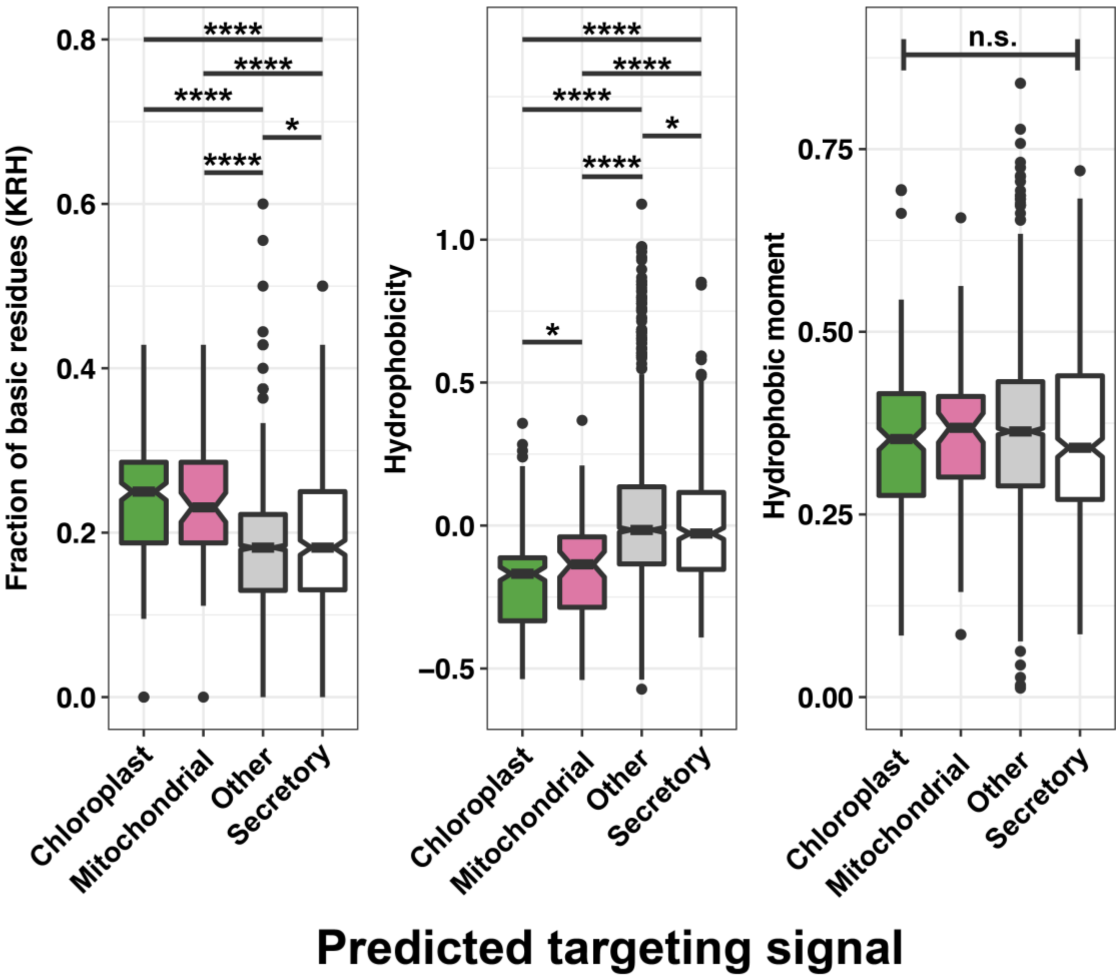


**Extended Data Fig. 30** Amphipathic elements in HeLEA1 homologs carrying different predicted targeting signal share different compositions in basic residues and hydrophobicity, but with same hydrophobic moment. (*: p<0.05, ****: p<0.0001, n.s.: not significant across all entries, Kruskal-Wallis rank sum test). Mitochondrial and chloroplast homologs tend to have higher amount of basic residues and less hydrophobic, indicating the lipid binding is mediated more through electrostatic interactions rather than hydrophobic interactions. The amphipathic elements are identified using the algorithm described in **Extended Data Fig. 11.** Each box represents the IQR of the dataset, whiskers represent plus/minus 1.5 IQR from the box hinge, and outliers are plotted as dot. Sample size: Chloropast: 220, Mitochondrial: 75, Secretory: 447, other: 2515.
